# Rotational settings quantize nucleosome movement by chromatin regulators

**DOI:** 10.64898/2025.12.12.693854

**Authors:** Van La, Abby Trouth, Vijay Ramani, Srinivas Ramachandran

## Abstract

Proper nucleosome positioning is essential for gene regulation and genomic integrity. Regulated nucleosome assembly and positioning results from a need to protect DNA sequences genome-wide, constrained by the known intrinsic sequence preferences of histones. Current models posit that chromatin regulators override the intrinsic preferences to establish the nucleosome landscapes observed *in vivo*, implying minimal roles for DNA sequence in guiding nucleosomal structure in cells. In contrast, we demonstrate that DNA sequence intrinsically guides the structure and remodeling of nucleosomes from yeast to mammals. We demonstrate that nucleosomes with weak translational settings *in vitro*, in yeast, and in mammalian cells demonstrate a clear preference for inward-facing A/T dinucleotides and outward facing G/C dinucleotides, at 10 bp spacings consistent with established preferences for rotationally positioned nucleosomes. Foreign DNA sequences heterologously inserted into the yeast genome obey similar rules, indicating that DNA sequence itself is causal. Finally, remodelers and transcription elongation change the preference among the alternative translational positions 10 bp apart, retaining the rotational setting. From these results, we propose that DNA sequence creates an energy landscape with preferred rotational settings every ∼10 bp, and that chromatin regulators, rather than overriding these preferences, navigate within them. This “detent” mechanism provides a unifying framework for understanding how diverse cellular processes achieve precise nucleosome positioning while maintaining the same DNA face exposed to regulatory factors.

## INTRODUCTION

Nucleosome distribution reflects genomic function.^1^ Active genes have a depletion of nucleosomes at the promoter and ordered nucleosomes upstream and downstream of the promoter.^2^ Inactive genes have significant nucleosome occupancy over their promoters.^3^ Nucleosomes are also ordered upstream and downstream of the binding sites of transcription factors (TF) like CTCF^4^, REST^5^, and PU.1.^6^ Loss of the three sliding remodelers (triple knockout or “TKO”), ISWI1, ISWI2, and Chd1 results in a dramatic loss of nucleosome periodicity and increased distance between nucleosomes at genes in budding yeast.^7^ In mouse embryonic stem cells (mESCs), loss of remodelers results in reduced accessibility at DNase I hypersensitive sites (DHSs) within minutes.^8^ Loss of the BPTF subunit or the ATPase subunit Snf2h (mammalian ISWI) of the NURF complex results in nucleosome gain at transcription start site (TSS)-distal DHSs with CTCF motifs.^9^ These drastic changes in the nucleosome landscape due to the loss of chromatin remodelers indicate that remodelers are essential for creating the observed strong nucleosome positioning patterns at active genes and TSS-distal DHSs.

In yeast, at least one of the three sliding remodelers is essential for yeast to survive stress conditions.^10^ The loss of nucleosome positioning in TKO leads to increased cryptic transcription and sensitivity to stress and genotoxic agents. In mESCs, the change in the nucleosome landscape associated with loss of BPTF is accompanied by impacts on gene expression, long-range chromatin interactions, and a loss of insulation at CTCF sites.^9^ In summary, maintaining proper nucleosome positions is essential for genome function.

Replication and transcription are some of the major causes of nucleosome shifts or disruptions, requiring reassembly and/or repositioning in the wake of replication^11^ and transcriptional machinery.^12^ Inhibition of transcription or the loss of elongation factors in yeast results in nucleosomes shifted downstream compared to their WT positions in genes. This molecular phenotype is rescued to an extent in the TKO background, indicating that transcription elongation and spacing remodelers move nucleosomes in opposite directions.^13–16^

Given the critical role of nucleosome positioning in gene regulation, long-range chromatin interactions, and protection against genotoxic stress, among other functions, determinants of nucleosome positioning have been heavily investigated.^17,18^ An overarching question driving these studies is the extent to which the underlying sequence drives nucleosome positioning. Sequence may influence nucleosome positioning directly, through effects on nucleosome stability, or indirectly, through transcription factor binding and sequence-dependent remodeler activity. The rotational and translational positions are the two parameters that describe nucleosome position (**Figure 1A**). Rotational position refers to the side of the DNA double helix in contact with the histone octamer^19^. Translational position refers to the base-pair location of the nucleosome dyad^19^. Nucleosome reconstitution studies on yeast genomic sequences *in vitro* have demonstrated that nucleosome depletion at promoters is supported by sequence to an extent^18,20^. However, translational positions around promoters that give rise to characteristic nucleosome arrays are not sequence-encoded^18,21,22^. Thus, the consensus is that remodelers override intrinsic sequence preferences to generate stereotypic nucleosome positioning genome-wide that supports genome function in context-specific ways.

**Figure 1.**
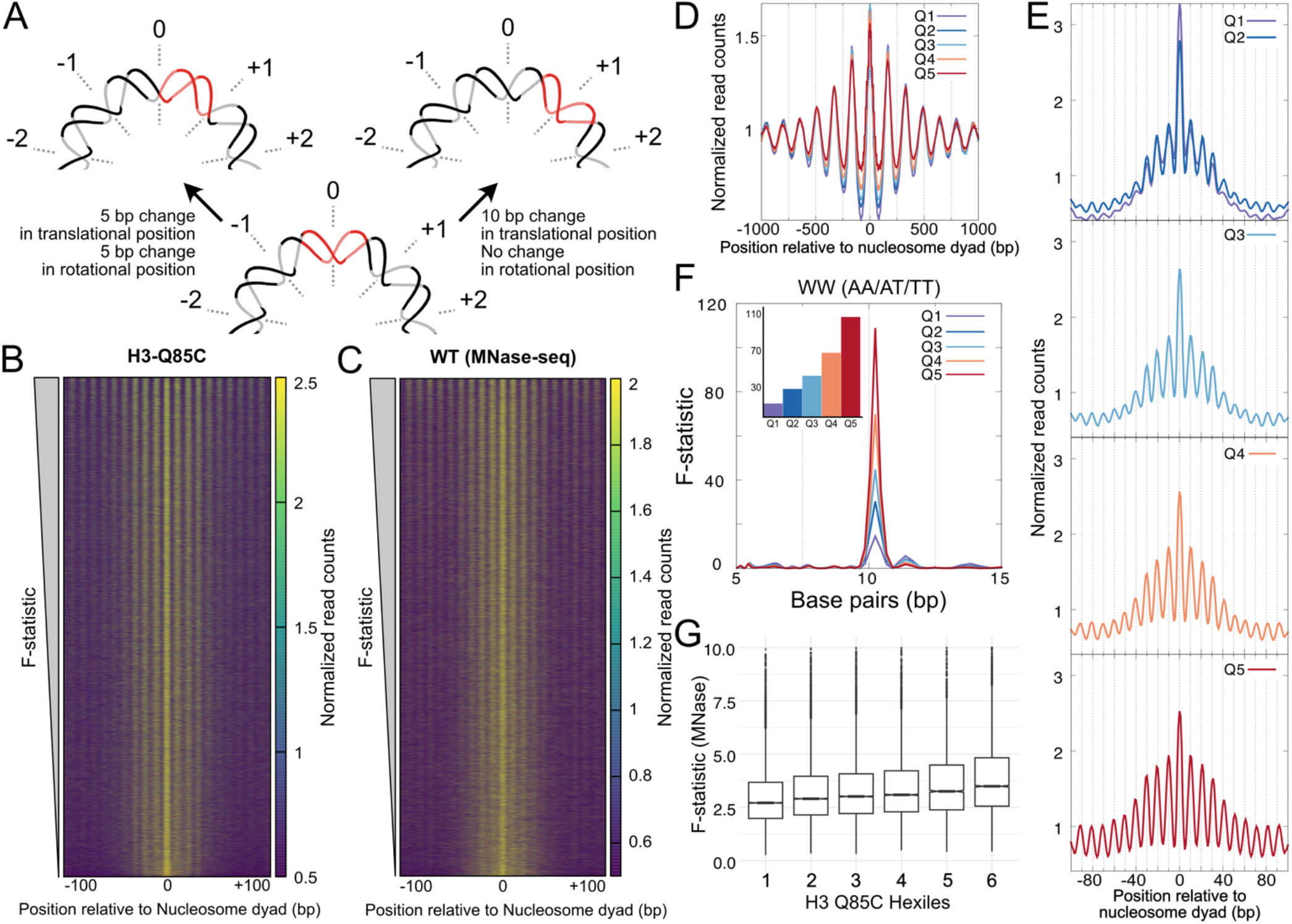
Preferred rotational settings observed at individual nucleosome positions. (**A**) Schematic showing effects of changes in rotational or translational positioning on the interactions between a specific 10 bp section of DNA and the histone octamer surface. 10 bp changes in translational position (which corresponds to no change in rotational position) results in no change in the face of the DNA segment interacting with histone octamer surface. Whereas a 5 bp change in translational position results in also a 5 bp change in rotational position, resulting in the change in the face of the DNA segment that interacts with the histone octamer. The dashed lines and the numbers denote the superhelical locations (SHL) with SHL-0 corresponding to the nucleosome dyad. (**B**) High-resolution coverage of fragment (50±2 bp) centers from H3-Q85C chemical cleavage mapping data at nucleosome positions across the genome, sorted by the strength of rotational setting (F-statistic from multitaper) plotted as a heatmap. (**C**) High-resolution coverage of fragment (147±5 bp) centers from MNase-seq data at nucleosome positions across the genome, sorted by the strength of rotational setting (F-statistic from multitaper) plotted as a heatmap. (**D**) Low-resolution coverage of fragments (50±2 bp) from H3Q85C data averaged across nucleosome positions divided into quintiles based on the strength of rotational setting (F-statistic from multitaper), plotted 1000 bp relative to nucleosome center. (**E**) High-resolution coverage of fragments (50±2 bp) from H3Q85C data averaged across nucleosome positions divided into quintiles based on the strength of rotational setting (F-statistic from multitaper), plotted 100 bp relative to nucleosome center. (**F**) Statistical measure of periodicity measured by multitaper of WW dinucleotide frequency for nucleosome positions in each quintile shows a strong peak around 10 bp that is higher for higher quintiles. Inset shows a bar plot for the value of the F-statistic at the peak around 10 bp. (**G**) Distribution of F-statistic reflecting strength of rotational setting (from multitaper) from MNase-seq data for nucleosome positions divided into hexiles shown as boxplots. Hexiles were defined based on the F-statistic calculated from H3-Q85C data. The boxplots show a positive correlation between the strengths of rotational settings observed with H3-Q85C data and that from MNase-seq.

It has long been appreciated that intrinsically flexible sequences drive rotational positioning. DNA fragments derived from MNase digestion of chicken erythrocytes showed a 10 bp periodicity of A/T (WW) and G/C (SS) dinucleotides, with WW dinucleotides facing inward and SS dinucleotides facing outward of the nucleosome.^23^ *In vitro* reconstitution of nucleosomes with yeast genomic DNA followed by MNase-seq also showed 10 bp WW dinucleotide periodicity, hinting at similar rules governing rotational positioning *in vitro* and *in vivo*.^24^ However, the fact that most translational positions are not similar *in vivo* versus *in vitro* has yet to be reconciled with the rules governing rotational positioning being the same *in vivo* and *in vitro*. Base-pair resolution is required to unambiguously assign rotational settings: a few bp variance does not have a significant impact on the interpretations derived from translational positions but can completely change the rotational setting of the nucleosome (**Figure 1A**).

Base-pair resolution nucleosome positions, made possible through chemical cleavage mapping, showed the extent of rotational phasing present in the yeast genome^25^ (and later in the mouse genome^26^). H4-S47C-anchored chemical cleavage mapping^25^ and later H3-Q85C-anchored chemical cleavage mapping^27^ showed that a consequence of strong rotational settings genome-wide is that each strong translational position from low-resolution mapping represents one of several translational positions 10 bp apart, which all share the same rotational setting. When nucleosome positions from chemical cleavage mapping are aligned genome-wide, the rotational settings show a strong 10 bp periodicity in WW and SS dinucleotides, with WW and SS dinucleotide frequency being anti-correlated. This dinucleotide periodicity is much stronger in nucleosome dyads identified from chemical cleavage mapping compared to those identified from MNase-seq datasets, possibly due to bp-resolution afforded by chemical cleavage mapping. Thus, chemical cleavage mapping has strengthened the observations supporting the early rules for the rotational phasing of nucleosomes.

While the existence of rotational positioning and its correlation with WW and SS dinucleotide periodicity has been appreciated for decades, several fundamental questions remain unanswered. First, can we quantify the strength of rotational positioning at individual nucleosome positions across the genome, rather than relying on genome-wide averages? Second, is the rotational positioning observed *in vivo* truly intrinsic to DNA sequence, or is it shaped by chromatin regulators? Third, and most critically, do cellular processes such as transcription and chromatin remodeling override or preserve these rotational settings? Answering these questions requires both high-resolution mapping methods and computational approaches capable of scoring rotational strength at single-nucleosome resolution.

Here, we address these questions through four advances. First, we develop a quantitative framework, based on multitaper spectral analysis, to score the strength of rotational setting at individual nucleosome positions genome-wide, rather than relying on genome-wide averages. Second, we show that this rotational information is recoverable from MNase-seq, the most widely used chromatin mapping method, greatly expanding the datasets to which rotational analysis can be applied. Third, using datasets^28^ generated from foreign DNA introduced into yeast as artificial chromosomes, we provide evidence consistent with rotational settings arising from sequence. Fourth, we systematically test whether these settings are maintained across perturbations of chromatin remodeling and transcription in yeast and mouse.

These experiments share a common logic: if chromatin regulators set rotational positions, then rotational settings should depend on the regulators rather than on the underlying sequence. Across every experiment we observed the opposite: rotational settings remained consistent with the sequence and did not change when regulators were removed or transcription was perturbed. We therefore conclude that chromatin regulators and active processes do not systematically override *in vivo* rotational settings but instead use the intrinsic sequence preferences of nucleosomes to mobilize them in 10 bp steps, changing only the relative preference among translational positions for a given rotational setting. Similar results in mESCs indicate that this is a general feature. In summary, many regions of the yeast and mouse genomes create a weak, sequence-based potential that guides nucleosome assembly and mobilization.

## RESULTS

### Preferred rotational settings observed at individual nucleosome positions

Chemical cleavage mapping in budding yeast confirmed preferred rotational settings for nucleosomes that correlate with ∼10 bp periodicity of WW dinucleotide frequency observed with earlier MNase-seq experiments, and also suggested alternative translational positions with the same rotational settings across the genome by averaging the chemical cleavage signal from all nucleosome positions^27^. Here, we first wanted to quantify the strength and prevalence of these alternative translational positions that share the same rotational settings at single nucleosome positions from Brogaard *et al.*^25^ Stepwise alternative translational positions with the same rotational settings will feature a ∼10 bp periodicity of the nucleosome signal (**Figure 1A**)^25,27^. To detect the presence and strength of 10 bp periodicity at individual nucleosome positions, we used the “multitaper” method inspired by its usage in ribosome profiling data where it is used to look for 3-bp periodicity that represents the ribosome positions at each successive codon (Ribotaper^29^). Multitaper uses a set of orthogonal window functions or tapers applied to discrete signals, or in our case, rotational positions and dinucleotide frequencies, before the Fourier transformation is computed^29^. Similar to Ribotaper for ribosome profiling data, we implemented multitaper to estimate the strength of 10 bp periodicity in datasets that report on nucleosome density. Multitaper also estimates an F-statistic at each frequency, which enables us to set a significance threshold to determine if a given nucleosome position has a strong rotational setting as revealed by strong alternative translational positions 10 bp apart. In other words, the F-statistic allows us to determine the statistical significance of the rotational setting at each translational position. To characterize the nucleosome positions identified to have higher F-statistic compared to positions with lower ones, we plotted the density of centers of 50±2 bp fragments generated from published H3Q85C-anchored chemical cleavage mapping (H3Q85C cleaves nucleosomal DNA and releases 51-bp fragments) at all nucleosome positions sorted by the multitaper F-statistic at frequencies around 0.1 bp^-1^, as a heatmap (**Figure 1B**). Given the high fraction of nucleosome positions with a strong F-statistic (**Supplementary Figure S1A**), we can conclude that most nucleosome positions have a strong rotational setting. In other words, at most nucleosome positions, nucleosomes sample alternate translational positions 10 bp apart, which would preserve their rotational setting.

We used the multitaper F-statistic at a frequency of around 0.1 bp^-1^ to bin nucleosome positions into quintiles, with the top 20% denoted as q5 and the bottom 20% as q1. We first plotted the H3-Q85C data +/-1000 bp relative to the nucleosome position and observed similar distances between nucleosome peaks across the quintiles, but higher signal between the peaks at higher quintiles (**Figure 1D**). At +/-100 bp relative to the nucleosome position, we can observe the central peak from **Figure 1D** is comprised of multiple translational positions with the same rotational setting, 10 bp apart (**Figure 1E**). In q1 compared to higher quintiles, the central peak is much higher compared to translational positions away from the central peak, but all quintiles had readily discernable peaks corresponding to alternative translational positions. In summary, we can observe strong rotational phasing at individual nucleosome positions across the genome, and we can quantify the strength of rotational phasing at individual nucleosome positions.

Chemical cleavage mapping is not performed routinely, but MNase-seq has been used to measure changes in nucleosome positioning upon various perturbations. So, we asked if preferred rotational settings and the corresponding alternate translational positions were also observed in MNase-seq data. We analyzed 147±5 bp fragments from published paired-end MNase-seq datasets^30^ with a smoothing window of 2 bp (referred to as “high-resolution” or high-res heareafter) to define MNase-derived nucleosomal protections precisely. Even though the MNase-seq data were not as robust as H3Q85C chemical cleavage data, we still observe a substantial fraction of nucleosome positions featuring alternate translational positions with the same rotational setting (**Figure 1C, Supplementary Figure S1A**). Similar to the analysis of H3-Q85C data, we binned nucleosome positions into quintiles based on multitaper F-statistic at a frequency around 0.1 bp^-1^. To observe nucleosome organization at larger length scales, we plotted the combined density of a wide range of fragments similar to earlier studies (140-160 bp, smoothed with a 10 bp smoothing window, called “low-resolution” or low-res hereafter). The low-res plot for quintiles of nucleosome positions genome-wide from MNase-seq data was highly similar to that of H3-Q85C (**Supplementary Figure S2C)**. The high-resolution plot revealed individual translational positions sharing the same rotational setting, with alternative positions showing progressively higher occupancy in higher quintiles, a pattern consistent with H3-Q85C data (**Supplementary Figure S2D)**.

Previous studies have aligned sequences at nucleosome positions across the genome and observed a strong AA/AT/TT periodicity that matches positions of histone-DNA contacts.^17,23,25,31–35^ Here, we have the added advantage of scoring the significance of rotational setting for every individual nucleosome using the multitaper approach. We next calculated grouped dinucleotide frequencies (WW: AA/AT/TT and SS:GG/GC/CC) for all nucleosome sequences for each quintile. To quantify the strength of dinucleotide oscillations, we again used multitaper, but this time on the WW or SS oscillations. When we plot the F-statistic against base-pairs (representing 1/frequency from mulltitaper output), we observe a clear peak at around 10 bp, reflecting the strong 10 bp periodicity in WW oscillations (**Figure 1F**). Furthermore, the strong cluster (q5) has much higher F-statistic compared to the weak cluster (q1), indicating that stronger rotational settings feature stronger 10-bp WW oscillations. This is true in both MNase-seq (**Supplementary Figure S2A**) and Q85C chemical cleavage (**Figure 1F**) datasets, strongly suggesting that this is an intrinsic feature of the sequence rather than being imposed by experimental variables. Furthermore, the F-statistic from MNase-seq data increases on average with increasing quintiles defined using H3-Q85C dataset, showing a concordance in strength of rotational settings observed by the two methods (**Figure 1G**).

To compare the strength of rotational setting at individual nucleosome positions across datasets and to dinucleotide frequencies of the underlying sequence, we developed a quantitative measure of the strength of rotational setting at any given translational position. We first averaged the nucleosome dyad frequencies of the top quintile of nucleosome positions based on multitaper F-statistic. We then identified peaks and troughs for the dyad distribution, which allowed us to calculate the Fractional Enrichment at Rotational Positions (FERP) score for each nucleosome position (**Supplementary Figure S1B**). FERP is the average of the difference in nucleosome occupancy at the peaks relative to the adjacent troughs, reflecting how well nucleosomes adopt the same rotational setting at any given locus. The FERP score is positive when nucleosome coverage is enriched at the expected rotational peak positions relative to flanking troughs, indicating a strong rotational setting at the locus. Since FERP scores are derived with internal normalization at each locus, they are well suited for quantitative comparison across datasets. We observe a strong and significant correlation of FERP scores derived at ∼67,000 nucleosome positions using the H3-Q85C dataset and the MNase-seq dataset, independently validating the concordance in strength of rotational settings observed between the two methods (**Supplementary Figure S1D**). We next developed a similar metric for dinucleotide frequencies. Each base-pair position in the nucleosome window was assigned a dinucleotide score of +1 (WW: AA, TT, AT, TA), -1 (SS: GG, CC, GC, CG), or 0 (mixed). We then identified peaks and troughs for the WW dinucleotide frequency of the top quintile of nucleosome positions based on multitaper F-statistic (**Supplementary Figure S1C**), and calculated the dinucleotide enrichment at rotational positions (DERP), similar to FERP, but in sequence space. We observed significant positive correlation between DERP and FERP for both H3-Q85C and MNase-seq datasets, confirming at the level of individual nucleosome positions, stronger rotational phasing correlates with stronger WW/SS dinucleotide oscillations (**Supplementary Figure S1K, L**). Thus, the high-resolution nucleosome positions from chemical cleavage mapping combined with analysis of 147±5 bp fragments from MNase-seq allows us to map rotational settings and alternative translational positions of nucleosomes using MNase-seq data.

### Yeast rotational positions do not arise from MNase bias

The strong rotational settings we observed could potentially arise from MNase sequence bias rather than true nucleosome positioning. Rotational settings are obvious in both H4-S47C and H3-Q85C cleavage mapping datasets, strongly suggesting that they do not arise from MNase sequence bias. However, to directly rule out digestion bias, we analyzed published sequencing datasets generated by digesting naked yeast genomic DNA with MNase^36^. With the low-res analysis, we do not see significant translational nucleosome positions (**Supplementary Figure S2E**). With high-res analysis, we do not see any signal similar to rotational settings observed *in vivo* (**Supplementary Figure S2F**). The minor peaks we do observe do not overlap *in vivo* translational positions that are at 10 bp multiples from the dyad position. Consistent with lack of rotational phasing, we observe negligible correlation of FERP scores of the naked DNA dataset with that of H3-Q85C and MNase-seq datasets (**Supplementary Figure S1E, F**).

To further rule out MNase sequence bias, we grouped nucleosome positions based on quintiles of F-statistic from multitaper analysis of MNase-seq dataset generated from digestion of naked DNA. Whereas, the yeast chromatin MNase-seq dataset shows a clear trend of higher quintiles of rotational settings having stronger WW dinucleotide frequency (**Supplementary Figure S2A**), digestion of naked DNA did not show differences between quintiles (**Supplementary Figure S2B**). Furthermore, DERP scores have negligible correlation with naked DNA FERP scores (**Supplementary Figure S1M**). Based on these analyses, we conclude that the rotational positions that we have observed are not an artifact of MNase sequence bias.

### Rotational settings observed *in vivo* reflect intrinsic nucleosome preferences

Having established that rotational settings are prevalent and quantifiable *in vivo*, we next asked whether these settings reflect intrinsic sequence properties or require cellular machinery for their establishment. Decades of work on nucleosome reconstitutions suggest that the rotational settings might reflect the intrinsic sequence preferences of the histone octamer. To test this hypothesis with the ability to map the strength of rotational settings of individual nucleosome positions, we turned to datasets generated by reconstitution of nucleosomes on yeast genomic DNA in the absence of any remodelers or chromatin regulators. Chacin *et al.*^37^ used bacterial plasmids containing inserts of large segments of the yeast genome, which were mixed with recombinant histones at high salt. The authors investigated whether the origin-flanking DNA sequences intrinsically affected nucleosome positions and found that there was no intrinsic formation of phased arrays observed with the *in vitro* reconstitution using SGD (Salt Gradient Dialysis)^37^. In SGD, as salt concentration is reduced, the nucleosomes are formed based on their intrinsic DNA preference^37^. With the SGD dataset, we calculated the strength of rotational setting at each mappable nucleosome position (see **Methods**) and grouped the positions into quintiles. With a low-res plot, we could still observe a weak peak across quintiles at nucleosome positions defined *in vivo* (**Figure 2A**, compared to **Figure 1D**). Low-res mapping did not capture any other positions. However, rotational settings identical to *in vivo* settings are apparent when we plot centers of 147±5 bp (called “high-resolution” or high-res hereafter), with increasing nucleosome density at translational positions farther away from the central peak for higher quintiles (**Figure 2B**). The strong rotational setting and the concomitant alternative translational positions are apparent at individual nucleosome positions, similar to *in vivo* MNase-seq and H3-Q85C mapping (**Figure 2C**). We also observe a significant positive correlation in FERP scores at individual nucleosome positions (calculated at ∼21,670 nucleosome positions with SGD data) between both SGD and MNase-seq, and SGD and H3-Q85C datasets (**Supplementary Figure S1G, H**). Notably, the correlation of FERP scores between SGD and naked DNA datasets was negligible (**Supplementary Figure S1I**). The correlation matrix of FERP scores across H3-Q85C, MNase-seq, SGD, and naked DNA shows clear clustering of H3-Q85C, MNase-seq, and SGD, and naked DNA being the outlier (**Supplementary Figure S1J**). This further confirms at the individual nucleosome level that the rotational settings are similar across methods and conditions. Finally, higher quintiles of strength of rotational settings had stronger 10-bp periodicity of the WW dinucleotide pattern (**Figure 2D**) and we observed significant positive correlation between SGD FERP scores and sequence-based DERP scores across the ∼21,670 nucleosome positions. In summary, yeast genomic DNA inherently sets rotational settings for nucleosomes that are identical to those observed *in vivo*.

**Figure 2.**
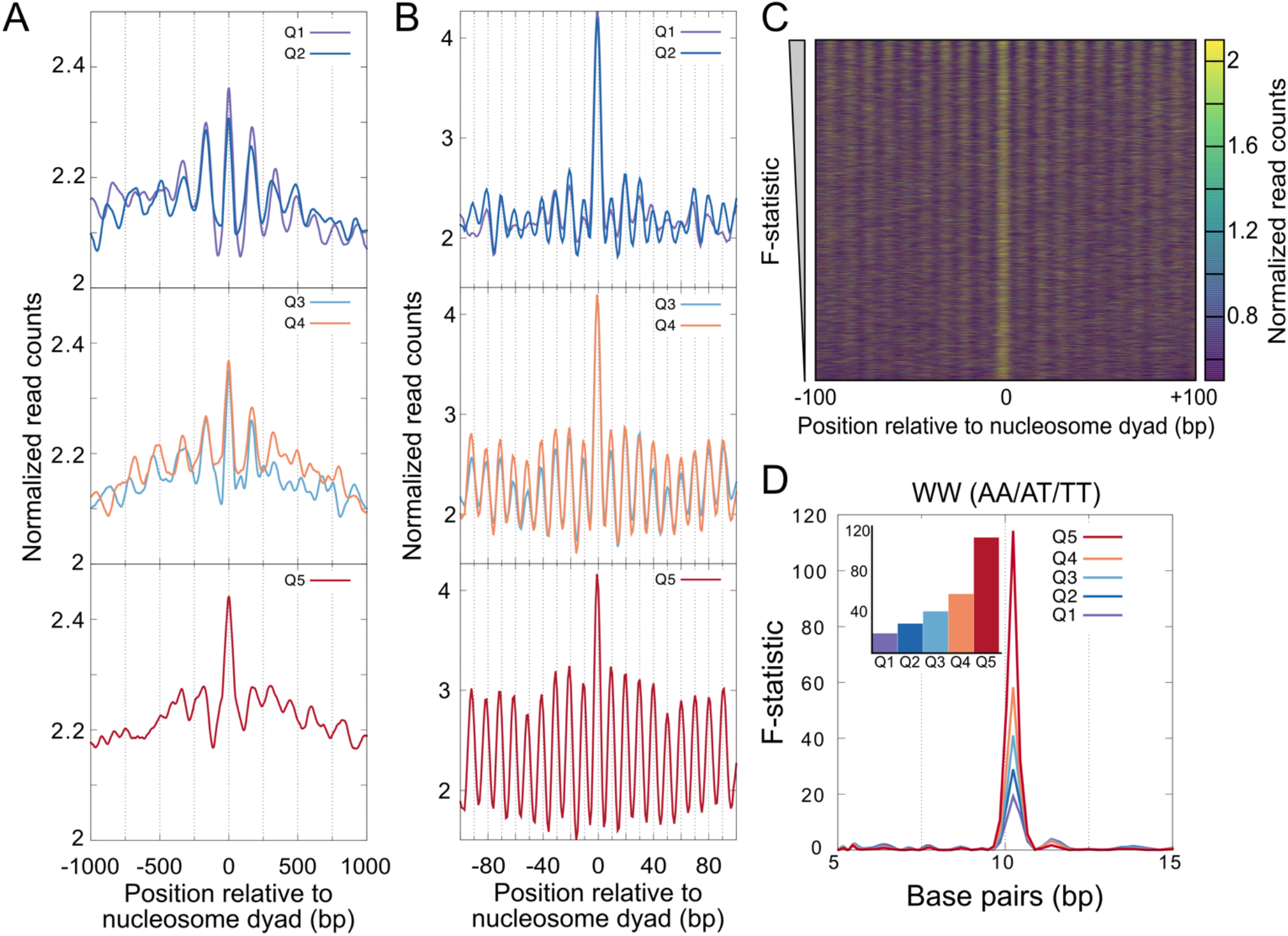
Rotational settings observed *in vivo* reflect intrinsic nucleosome preferences. (**A**) Low-resolution plot of MNase-seq data derived from nucleosomes reconstituted *in vitro* using SGD, for quintiles of nucleosome positions defined based on the strength of rotational phasing. Q1 represents the bottom 20% and Q5 represents the top 20% of nucleosome positions scored by their F-statistic from multitaper. (**B**) High-resolution plot of MNase-seq data derived from nucleosomes reconstituted *in vitro* using SGD for same quintiles as (**A**). (**C**) High-resolution coverage of fragment (147±5 bp) centers from SGD MNase-seq data at nucleosome positions across the genome, sorted by the strength of rotational setting (F-statistic from multtitaper) plotted as a heatmap. (**D**) Statistical measure of periodicity measured by multitaper of WW dinucleotide frequency for nucleosome positions in each quintile shows a strong peak around 10 bp that is higher for higher quintiles. Inset shows a bar plot for the value of the F-statistic at the peak around 10 bp.

### Intrinsically preferred rotational settings preserved by remodelers *in vitro*

The identical rotational settings observed in SGD and *in vivo* nucleosomes raised a key question: do chromatin remodelers override these intrinsic preferences when establishing functional chromatin architecture, or do they work within the constraints imposed by sequence? Chacin *et al.*^37^ in addition to reconstituting nucleosomes by SGD, also added purified remodelers to the reconstitutions and then mapped nucleosomes using MNase-seq. The low-res plots of datasets generated after addition of Chd1, INO80, RSC, ISW1a, ISWR1, and SWI/SNF show increased nucleosome occupancies, suggestive of increasing nucleosome density due to the action of remodelers (**Figure 3A-F**).

**Figure 3.**
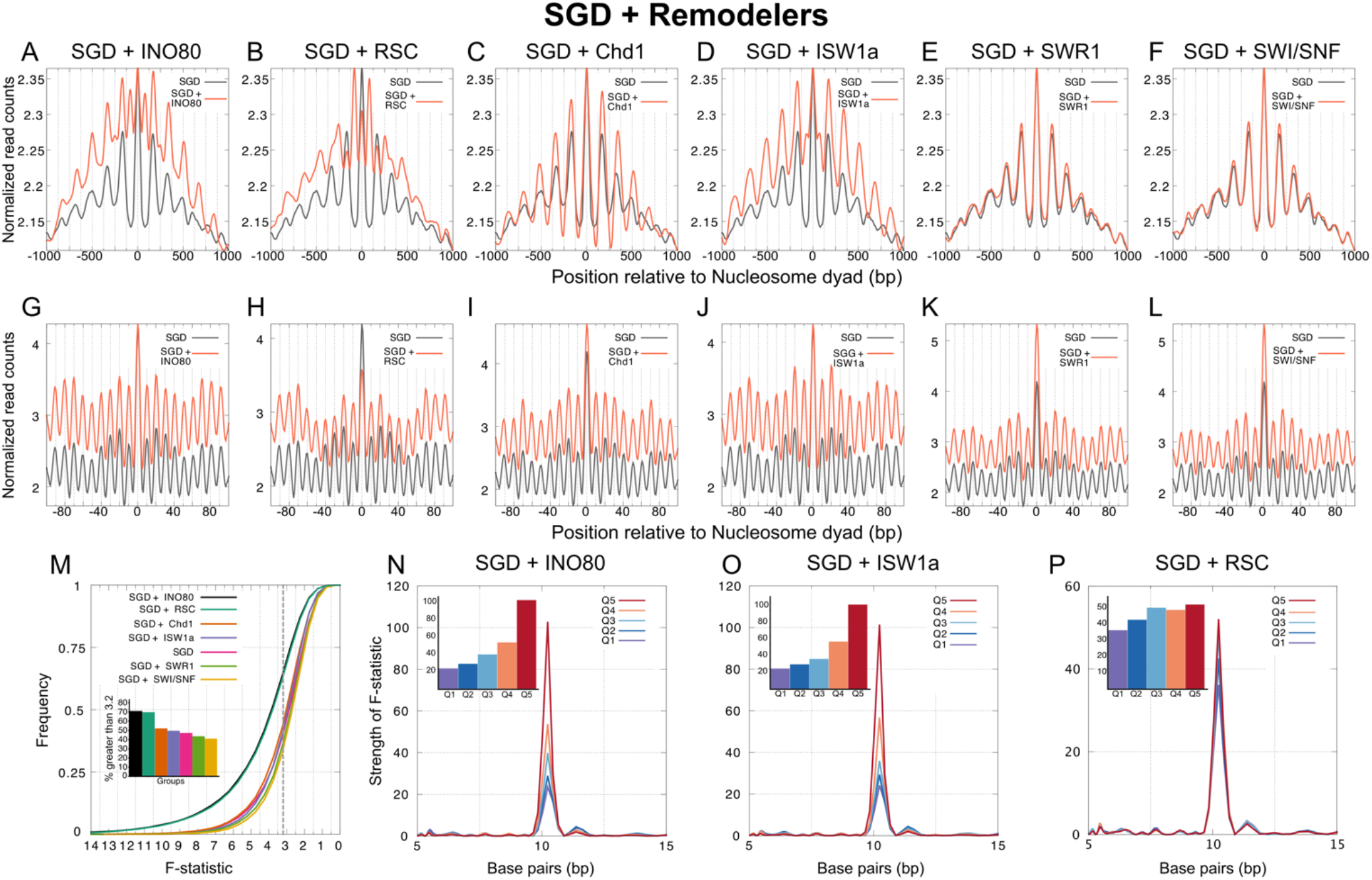
Intrinsically preferred rotational settings preserved by remodelers *in vitro*. (**A-F**) Low-resolution plot of MNase-seq data derived from nucleosomes reconstituted *in vitro* using SGD compared to SGD+remodelers at all nucleosome positions with coverage. (**G-L**) High-resolution plot of MNase-seq data derived from nucleosomes reconstituted *in vitro* using SGD compared to SGD+remodelers at all nucleosome positions with coverage. (**M**) Cumulative frequency of fraction of nucleosome positions as a function of decreasing F-statistic from multitaper for all nucleosome positions with plasmid coverage in SGD compared to SGD+remodelers. Dashed vertical line in gray is positioned at 3.2 (statistically significant, p-value < 0.05). The inset shows a bar-plot with % of nucleosome positions with F-statistic > 3.2. (**N-P**) Statistical measure of periodicity measured by multitaper of WW dinucleotide frequency for nucleosome positions in each quintile shows a strong peak around 10 bp that is higher for higher quintiles. Inset shows a bar plot for the value of the F-statistic at the peak around 10 bp.

In high-res plots, adding remodelers increased nucleosome signal, indicative of higher density, but did not shift peak positions relative to the SGD sample (**Figure 3G-L**). Remodelers INO80, RSC, Chd1, and ISW1a, when added to SGD nucleosomes, increase the fraction of positions genome-wide that are strongly rotational phased, whereas SWR1 and SWI/SNF slightly decrease the fraction of positions genome-wide that are strongly rotational phased (**Figure 3M**). For each of the remodeler datasets, we split nucleosome positions into quintiles based on the strength of rotational phasing and calculated the strength of WW dinucleotide frequency across sequences in each quintile. For all remodelers, as with SGD alone, WW dinucleotide periodicity strength increased with stronger rotational phasing (**Figure 3N-P, Supplementary Figure S3**). Thus, *in vitro*, remodelers increase nucleosome density while maintaining rotational settings of the nucleosomes, suggesting that remodelers take advantage of the 10 bp energy potential provided by the sequence but do not override it.

### Strength of rotational positioning correlates with the strength of WW periodicity in foreign DNA inserted into yeast genome

Our SGD analysis is consistent with yeast genomic DNA intrinsically encoding rotational settings and that the strength of WW dinucleotide periodicity correlates with strength of rotational positioning. However, yeast sequences have co-evolved with the histone octamer over millions of years. To test whether rotational positioning truly emerges from general sequence properties rather than yeast-specific features, we turned to an elegant system: foreign DNA sequences from diverse species introduced into yeast as artificial chromosomes. When foreign DNA is inserted to the yeast genome, Meneu *et al*. found that chromatin spontaneously forms even though these sequences have not co-evolved with the yeast’s regulatory machinery^28^. Meneu *et al*. further identified the sequence composition (GC content) as a primary determinant of transcriptional activity and state of chromatin.

The yeast strains with extra chromosomes made of foreign DNA offered an ideal system to ask if rotational settings are set by sequence. Using the published MNase-seq datasets from the yeast hybrid strains, we asked if there are preferred rotational settings in the foreign chromosomes. With 4 different strain combinations (Yeast + Mmyco (DNA from *Mycoplasma mycoides*), Yeast + Mpneumo (DNA from *Mycoplasma pneumoniae*), Yeast + Vitis (DNA from *Candidatus Phytoplasma vitis*), and Yeast + Falciparum (DNA from *Plasmodium falciparum*)), we first identified high-resolution nucleosome positions (see ***Methods***) and then generated low-res plots centered at the called nucleosome positions (**Figure 4A-D**). Similar to the original study, we observed nucleosome arrays of various sizes at the artificial chromosome from the four hybrid strains. The Mmyco, Mpneumo, and Vitis yeast hybrid strains feature artificial chromosome with sequences derived from a bacterial background while Falciparum yeast hybrid strain’s artificial chromosome is from a eukaryotic background. When we generated the high-res plots centered at all called nucleosome positions of the artificial chromosomes, we observed 10 bp periodicity in nucleosome density for Mmyco and Vitis sequences, but not for Mpneumo and Falciparum sequences, although all four show highest nucleosome density at the central position (**Figure 4E-H**). We then scored the strength of rotational settings at each position of the artificial chromosomes and as suggested by the high-res plots, Mmyco sequences have the highest fraction of positions with significant rotational positioning, followed by Vitis, Falciparum, and Mpneumo respectively (**Figure 4I**). We then determined the strength of WW oscillations at nucleosome positions for sequences from each of these species. Strikingly, the fraction of nucleosome positions with significant rotational positioning correlated with the strength of 10-bp oscillations of the WW dinucleotide frequency, with Mmyco having the strongest WW oscillations, followed by Vitis, Mpneumo, and Falciparum respectively (**Figure 4J**). Thus, the nucleosome positioning in artificial chromosomes containing non-yeast sequences is also consistent with rotational settings being driven by WW dinucleotides every 10 bp. To further ask if WW dinucleotide frequency can explain the strength of rotational phasing within each artificial chromosome, we identified nucleosome positions with strongest (top 10% of positions) and weakest (bottom 10% of positions) rotational settings for the non-yeast sequences of each hybrid strain. The low-res analysis shows the same features in the foreign sequences that was observed in yeast (**Supplementary Figure S4A-D**) and the rotational settings become more apparent in the strong positions in the high-res analysis (**Supplementary Figure S4E-H**). Strikingly, even in the exogenous sequences with weak WW oscillations globally, the subset of positions with strong rotational phasing have stronger WW oscillations compared to positions with weak rotational phasing (**Supplementary Figure S4I, J**). The artificial chromosomes with the weakest rotational phasing did not show appreciable WW oscillations as expected (**Supplementary Figure S4K, L**). In summary, rotational settings encoded by WW oscillations are apparent even in sequences derived from species without nucleosomes when these are introduced as artificial chromosomes in yeast.

**Figure 4.**
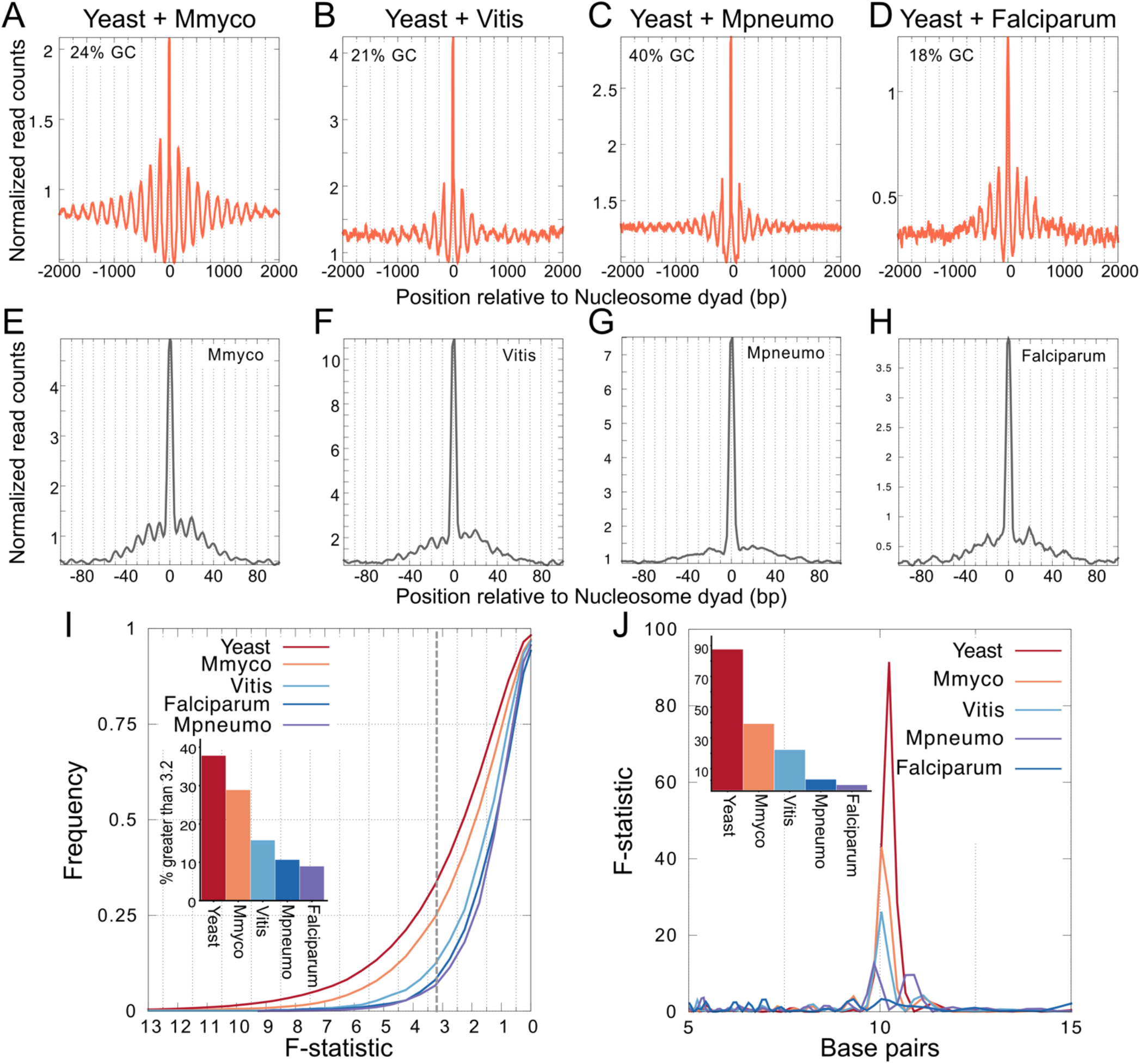
Strength of rotational positioning correlates with the strength of WW periodicity in foreign DNA inserted into yeast genome. (**A-D**) Low-resolution plot of MNase-seq data derived from yeast hybrid strains strain combinations: Yeast + Mmyco (DNA from *Mycoplasma mycoides*), Yeast + Vitis (DNA from *Candidatus Phytoplasma vitis*), Yeast + Mpneumo (DNA from *Mycoplasma pneumoniae*), and Yeast + Falciparum (DNA from *Plasmodium falciparum*), at all nucleosome positions identified on the exogenous DNA sequences. (**E-H**) High-resolution plot of MNase-seq data for same strains as (**A-D**). (**I**) Cumulative frequency of fraction of nucleosome positions as a function of decreasing F-statistic from multitaper for all nucleosome positions on the exogenous DNA sequences. The inset shows a bar-plot with % of nucleosome positions with F-statistic > 3.2. (**J**) Statistical measure of periodicity measured by multitaper of WW dinucleotide frequency for nucleosome positions in each exogenous sequence shows a strong peak around 10 bp for exogenous sequences that have strong rotational positioning. Inset shows a bar plot for the value of the F-statistic at the peak around 10 bp.

### Nucleosome shifts due to perturbations in remodeling are quantized

Our *in vitro* analyses established that rotational settings are sequence-intrinsic and preserved by remodelers acting on reconstituted chromatin. We next asked whether this principle holds *in vivo*, where multiple remodelers, transcription, and other cellular processes continuously mobilize nucleosomes. If rotational settings truly constrain nucleosome dynamics, they should persist even when these processes are perturbed. To determine the role of chromatin remodelers in driving the observed rotational settings for nucleosomes *in vivo*, we first analyzed MNase-seq datasets^13^ from yeast strains containing deletions of three chromatin remodelers, ISWI1/2, CHD1 (triple knockout, “TKO”). Singh *et al*.^13^ reported cryptic transcription when nucleosome arrays were disrupted due to the absence of these remodelers in the TKO strain.^13^ Based on low-res plots, it has been observed that TKO displays a loss of nucleosome periodicity upstream and downstream of promoters^13^ (**Supplementary Figure S5A**). For example, at the +4 position downstream of the TSS, the effect of TKO is stark at low-res: there is a complete loss of positioning (**Supplementary Figure S5B**). Genome-wide, TKO results in drastic loss of nucleosome density at WT nucleosome positions and increased distances between nucleosome peaks (**Figure 5A, B**).

**Figure 5.**
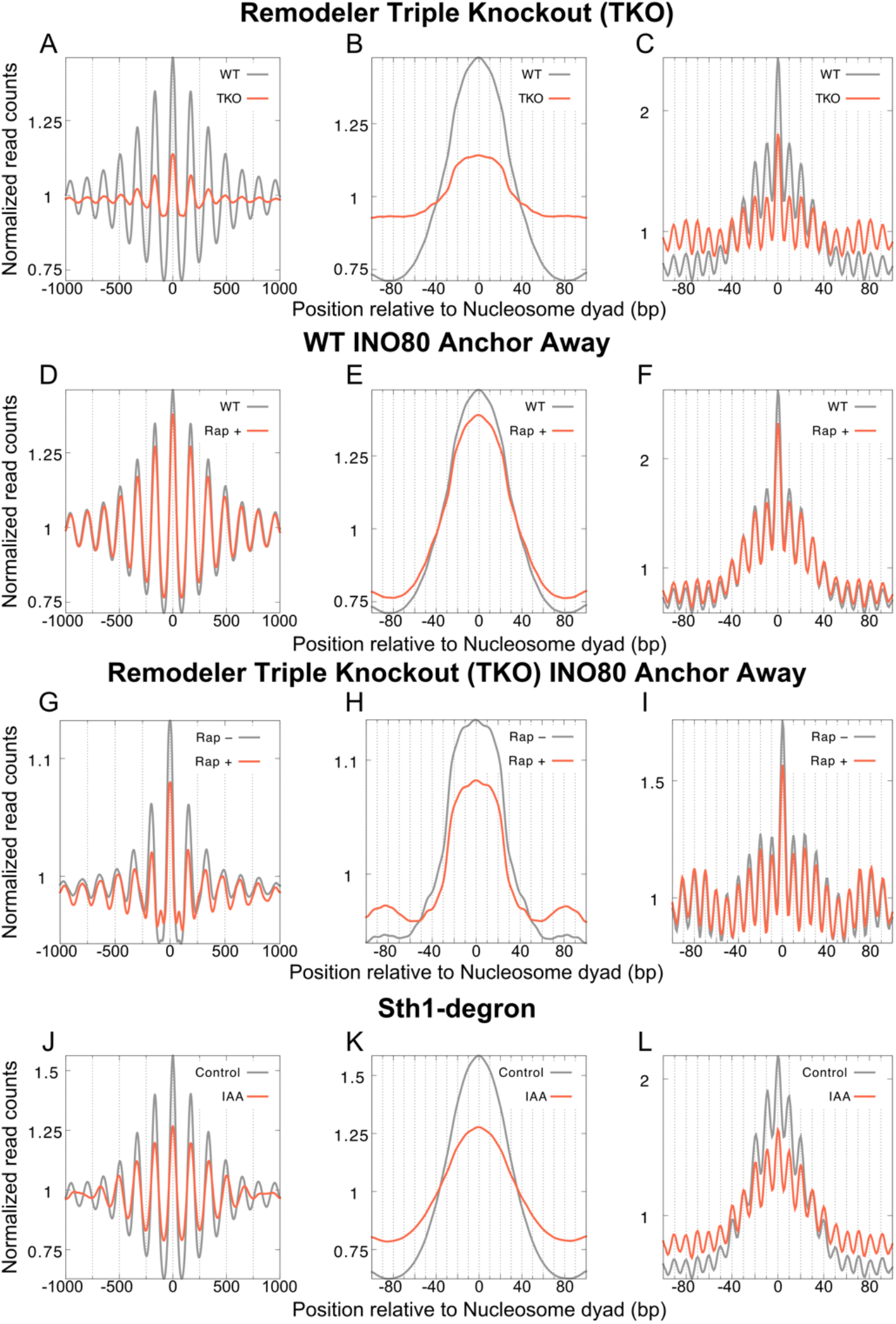
Nucleosome shifts due to perturbations in remodeling are quantized. (**A**) Low-resolution plot of MNase-seq data derived from TKO strain versus WT at +/-1000 bp relative to nucleosome positions genome-wide. (**B**) Low-resolution plot of MNase-seq data derived from TKO strain versus WT at +/-100 bp relative to nucleosome positions genome-wide. (**C**) High-resolution plot of MNase-seq data derived from TKO strain versus WT at +/-100 bp relative to nucleosome positions genome-wide. (**D-F**) Same as (**A-C**) for INO80 anchor-away strain with rapamycin treatment for 90 minutes compared to no rapamycin. (**G-I**) Same as (**A-C**) for TKO/INO80 anchor-away strain with rapamycin treatment for 90 minutes compared to no rapamycin. (**J-L**) Same as (**A-C**) for strain containing Sth1 tagged with auxin-inducible degron with indole-3-acetic acid (IAA) treatment for 60 minutes compared to no IAA treatment.

Strikingly, with high-res plots, we observe the rotational setting is maintained in the TKO strain at all nucleosome positions genome-wide (**Figure 5C**) and in the gene bodies as well (**Supplementary Figure S5C, D**). The preferred translational position in TKO is still the central position; however, the alternate translational positions upstream and downstream of the preferred position gain in occupancy. We next asked if this trend holds true at individual nucleosome positions. We compared the FERP scores across all nucleosome positions from WT and TKO datasets and observed a strong positive correlation (r=0.79, p<2.2×10^-16^), suggesting that even at individual nucleosome positions, rotational settings are conserved upon loss of the three positioning remodelers (**Supplementary Figure S6A**). To ask whether any individual position deviates from the WT-TKO relationship more than expected, we externally studentized the residuals of the linear fit and found no residual with an adjusted p-value below 0.05. This suggests that at the level of detection afforded by MNase-seq, there are no nucleosome positions that significantly deviate in strength of rotational setting when comparing WT and TKO strains. As a second, independent check that TKO preserves rotational settings, we compared WT and TKO at the level of individual replicates: the correlation between WT and TKO replicates (R = 0.66-0.70) is similar to the correlation between replicates of the same genotype, with TKO setting the floor (TKO vs. TKO, R = 0.66; WT vs. WT, R = 0.76), indicating that genotype identity does not structure the FERP relationship beyond what replicate noise already accounts for (**Supplementary Figure S6B-G**). Thus, the loss of the three positioning remodelers, which results in a global loss of translational positioning, does not affect the rotational settings of nucleosomes. While remodelers may move nucleosomes in smaller increments, our analysis suggests that the stable positions observed at steady state are quantized in ∼10 bp intervals that preserve rotational settings, analogous to ‘rest stops’ along a continuous energy landscape.

The INO80 complex plays important roles in DNA repair, replication, and transcription. The INO80 complex is essential to position the +1 nucleosome *in vitro* and *in vivo*^38–40^. In addition, loss of INO80 affects nucleosome spacing in the gene body as well. Here we determined the effect of the loss of INO80 on rotational settings of genome-wide and in gene body nucleosomes using published data^13^. The low-res mapping of WT and TKO cells with depletion of INO80 shows mild loss of nucleosome density at WT nucleosome positions and slight increase in fuzziness of the nucleosome (**Figures 5D, E, G, H Supplementary Figure S5E, F, I, J**). However, with the high-res mapping, we were able to observe rotational settings intact genome-wide (**Figure 5F, I**) and in promoters and gene bodies (**Supplementary Figure S5G, H, K, L**).

We next analyzed nucleosome positioning upon the depletion of Sth1, the catalytic subunit of the RSC complex using published data^41^. RSC is essential for the formation of the NDR by keeping the -1 and +1 nucleosomes from encroaching into the NDR^40–43^. Here, we see this shift of nucleosomes into the NDR and shift of gene body nucleosomes towards the promoter in the low-res plots, as seen before (**Supplementary Figure S5M, N**). Genome-wide, we see a loss of nucleosome density at WT positions and increased distances between nucleosome peaks (**Figure 5J, K**). In high-res plots, it is apparent that the rotational settings do not change, and there is a clear gain in occupancy for translational positions upstream of the preferred dyad position of the WT and loss of occupancy of translational positions downstream of the preferred dyad position of the WT (**Figure 5L, Supplementary Figure S5O, P).** Taken together, we conclude that the observed rotational settings *in vivo* are not set by ISWI1/2 and CHD1 or INO80, or RSC alone. These remodelers change translational positions of nucleosomes without changing their rotational setting *in vivo*. Since rotational settings remain unchanged due to loss of remodelers, our results suggest that all nucleosomal shifts due to chromatin remodeling *in vivo* are quantized at 10 bp.

### Nucleosome shifts due to perturbations in transcription are also quantized

We next asked if rotational settings change upon perturbation of transcription. Transcription elongation across nucleosomes is thought to be akin to a Brownian ratchet where the movement of RNA polymerase II (RNAPII) sequentially unwraps DNA from the nucleosome directly upstream of the elongation complex.^44–46^ When we analyze MNase-seq data generated after the depletion of the major subunit of RNAPII, Rpb1^13^, we observe a downstream shift in nucleosome positions at low-res especially with increasing distance from the promoter, reproducing published results (**Figure 6A**, **6B**).^13,47^ However, in the high-res analysis, the alternate translational positions are apparent even after loss of Rpb1, pointing to rotational settings being the same regardless of transcription status (**Figure 6C**, **6D, Supplementary Figure S7A-C, Supplementary Figure S8A-C**).

**Figure 6.**
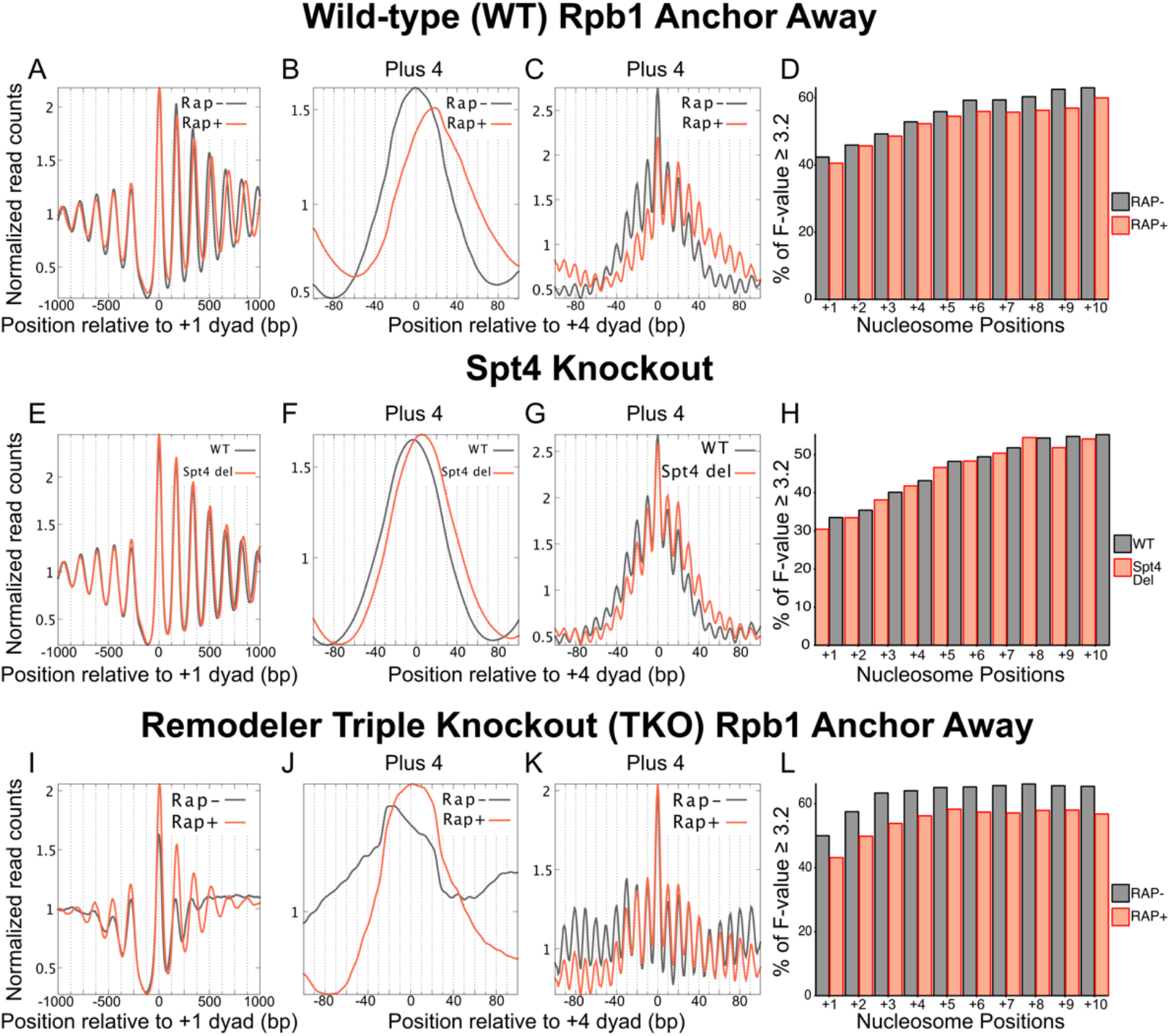
Nucleosome shifts due to perturbations in transcription are also quantized. (**A**) Low-resolution plot of MNase-seq data derived from Rpb1 anchor-away strain with rapamycin treatment for 120 minutes compared to no rapamycin at +/-1000 bp relative to the +1 nucleosome position across all genes. (**B**) Low-resolution plot of MNase-seq data derived from Rpb1 anchor-away strain with rapamycin treatment for 120 minutes compared to no rapamycin at +/-100 bp relative to the +4 nucleosome position across all genes. (**C**) High-resolution plot of MNase-seq data derived from Rpb1 anchor-away strain with rapamycin treatment for 120 minutes compared to no rapamycin at +/-100 bp relative to the +4 nucleosome position across all genes. (**D**) Fraction of genes with significant rotational phasing (F-statistic > 3.2) at each nucleosome position downstream of the TSS based on the MNase-seq data derived from Rpb1 anchor-away strain with rapamycin treatment for 120 minutes compared to no rapamycin. (**E-H**) Same as (**A-D**) comparing Spt4 knockout strain to WT. (**I-L**) Same as (**A-D**) for Rpb1 anchor-away strain in the TKO background with rapamycin treatment for 120 minutes compared to no rapamycin.

Transcription elongation through chromatin is assisted by several factors including specific elongation factors. When an elongation factor such as Spt4 is removed, RNAPII elongation is inhibited. Previous studies showed that Spt4 helps promote movement of RNAPII through the nucleosomes of the gene body especially the +2 nucleosome and regulate early elongation dynamics^16,48–50^. We analyzed published MNase-seq data^16^ generated after deletion of Spt4 as an alternative to loss of Rpb1, to ask if slower elongation shifts nucleosomes from their preferred rotational settings. When we reanalyzed MNase-seq data generated after deletion of Spt4, at low-res, we see a downstream shift in nucleosome positions, reproducing published results (**Figure 6E**, **6F**). However, at high-resolution, we observe a quantized shift downstream that is similar to the effect upon removing RNAPII from the nucleus (**Figure 6G**, **6H, Supplementary Figure S7D-F, Supplementary Figure S8D-F**). Similar effects observed for both the loss of Rpb1 and Spt4 validate our conclusion that inhibition of transcription still preserves the rotational setting of nucleosomes while at the same time increasing occupancy at translational positions downstream of the positions preferred in WT.

Finally, we analyzed nucleosome positions upon loss of Rpb1 in the TKO strain, which was shown to partially restore translational positions^13^ (**Figure 6I**, **6J**), leading to the hypothesis that the opposite effects of transcription and remodeling result in the observed steady-state nucleosome landscape. Here also, we observe the rotational settings preserved between TKO and TKO with loss of Rpb1 (**Figure 6K, L, Supplementary Figure S7G-I, Supplementary Figure S8G-I**). In summary, either stopping or slowing down RNAPII elongation results in a downstream shift in preferred translational positions of nucleosomes while preserving their rotational settings. Thus, neither remodeling nor transcription set rotational settings across the genome in budding yeast, rather they take advantage of preferred rotational settings in the budding yeast genome to quantize nucleosome mobilization in 10 bp steps.

### Mammalian nucleosomes feature strong rotational settings that do not change upon loss of NURF

Budding yeast allows nucleosomes to assemble at intrinsically preferred rotational settings *in vivo* both in the yeast genome and on genomes of other species introduced into yeast. The intrinsic preference correlates with the strength of 10 bp WW periodicity. We next asked if our quantized shift model is generalizable beyond budding yeast. Mammalian nucleosome positions are not as well mapped as yeast due to the much larger genome size and difficulty in implementing chemical cleavage mapping due to tens of copies of histone genes. We focused on mouse embryonic stem cells (mESCs), where H4S47C chemical cleavage mapping has been performed^26^. We generated a deep MNase-seq dataset (∼1 billion sequenced fragments). We calculated the strength of rotational settings at nucleosome positions genome-wide defined by Voong *et al.*^26^ using chemical cleavage mapping. We then ordered nucleosome positions by the multitaper F-statistic and plotted density of 147±5 bp fragments from our deep MNase-seq dataset as a heatmap (**Figure 7A**). Genome-wide, we observe alternative translational positions 10 bp apart, indicative of strong rotational settings. We split all nucleosome positions into five quintiles based on the multitaper F-statistic and generated a low-res plot at +/-1000 bp relative to the nucleosome center (**Figure 7B**). Similar to yeast, the central position has higher read density for lower quintiles whereas regions between nucleosome positions have higher read densities for higher quintiles, consistent with higher occupancy at alternative translational positions for nucleosome positions belonging to higher quintiles. The alternative translational positions 10 bp apart up to 80 bp from the nucleosome center are apparent with the high-res plot, with alternate positions having higher read density for positions belonging to higher quintiles (**Figure 7C**). We next calculated the strength of WW periodicity for each quintile and observed increasing quintiles have stronger WW periodicity (**Figure 7D**). Genome-wide, ∼25% of nucleosome positions are significantly rotationally phased (**Figure 7E**).

**Figure 7.**
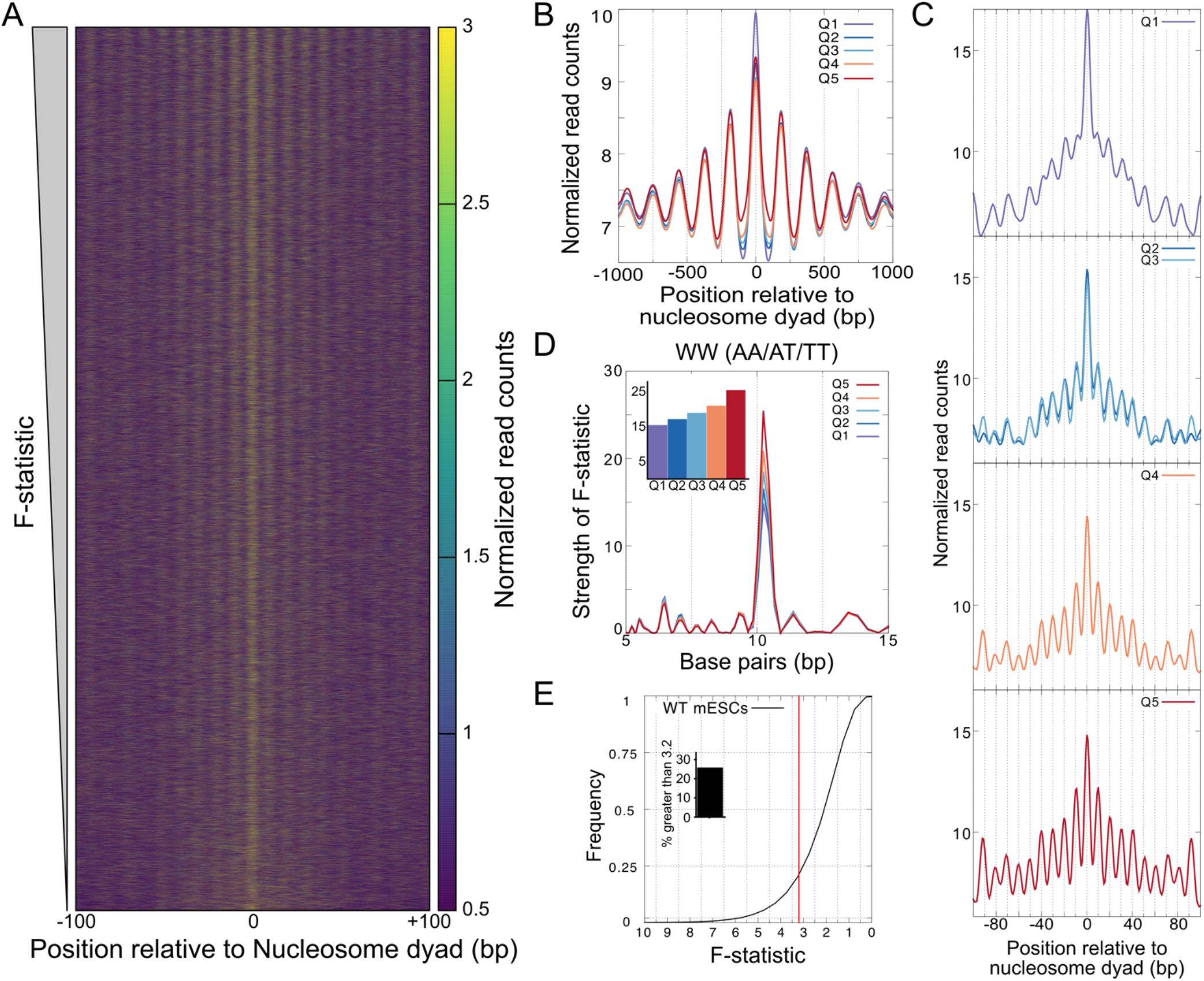
Mammalian nucleosomes feature strong rotational settings in mESCs. (**A**) High-resolution coverage of 147±5 bp fragment centers from MNase-seq performed with mESCs at nucleosome positions across the genome (defined from H4-S47C cleavage mapping data), sorted by the strength of rotational setting (F-statistic from multitaper) plotted as a heatmap. (**B**) Low-resolution plot of MNase-seq data from mESCs averaged across nucleosome positions divided into quintiles based on the strength of rotational setting (F-statistic from multitaper), plotted 1000 bp relative to nucleosome center. (**C**) High-resolution plot of MNase-seq data from mESCs averaged across nucleosome positions divided into quintiles based on the strength of rotational setting (F-statistic from multitaper), plotted 100 bp relative to nucleosome center. (**D**) Statistical measure of periodicity measured by multitaper of WW dinucleotide frequency for nucleosome positions in each quintile shows a strong peak around 10 bp that is higher for higher quintiles. Inset shows a bar plot for the value of the F-statistic at the peak around 10 bp. (**E**) Cumulative frequency of fraction of nucleosome positions as a function of decreasing F-statistic from multitaper for all nucleosome positions in the mouse genome. The inset shows a bar-plot with % of nucleosome positions with F-statistic > 3.2.

Next, to explore changes in rotational settings upon loss of remodelers, we focused on TSS-distal DNase-hypersensitive sites (DHS). Nucleosomes are depleted at the center of the DHS and previous studies have shown that loss of NURF component BPTF results in gain in nucleosomes at DHS centers and a relative shift of nucleosomes on either side of the DHS towards the DHS center^9^. To anchor our analysis at a nucleosome position, we chose the first nucleosome position downstream of the DHS (DHS+1). We identified the strongest and weakest 2000 DHS+1 nucleosome positions based on the strength of their rotational settings. The low-res plot shows similar nucleosome distribution relative to the DHS+1 nucleosome center for both sets of nucleosome positions, with slightly higher nucleosome depletion for the positions with weaker rotational phasing (**Figure 8A**). With high-res plot, the alternative translational positions 10 bp apart are apparent for positions with strong rotational phasing, whereas alternative translational positions are absent for the nucleosome positions with weak rotational phasing (**Figure 8B**). As observed genome-wide, the positions with strong rotational phasing have stronger WW periodicity at 10 bp compared to the weaker positions (**Figure 8C**).

**Figure 8.**
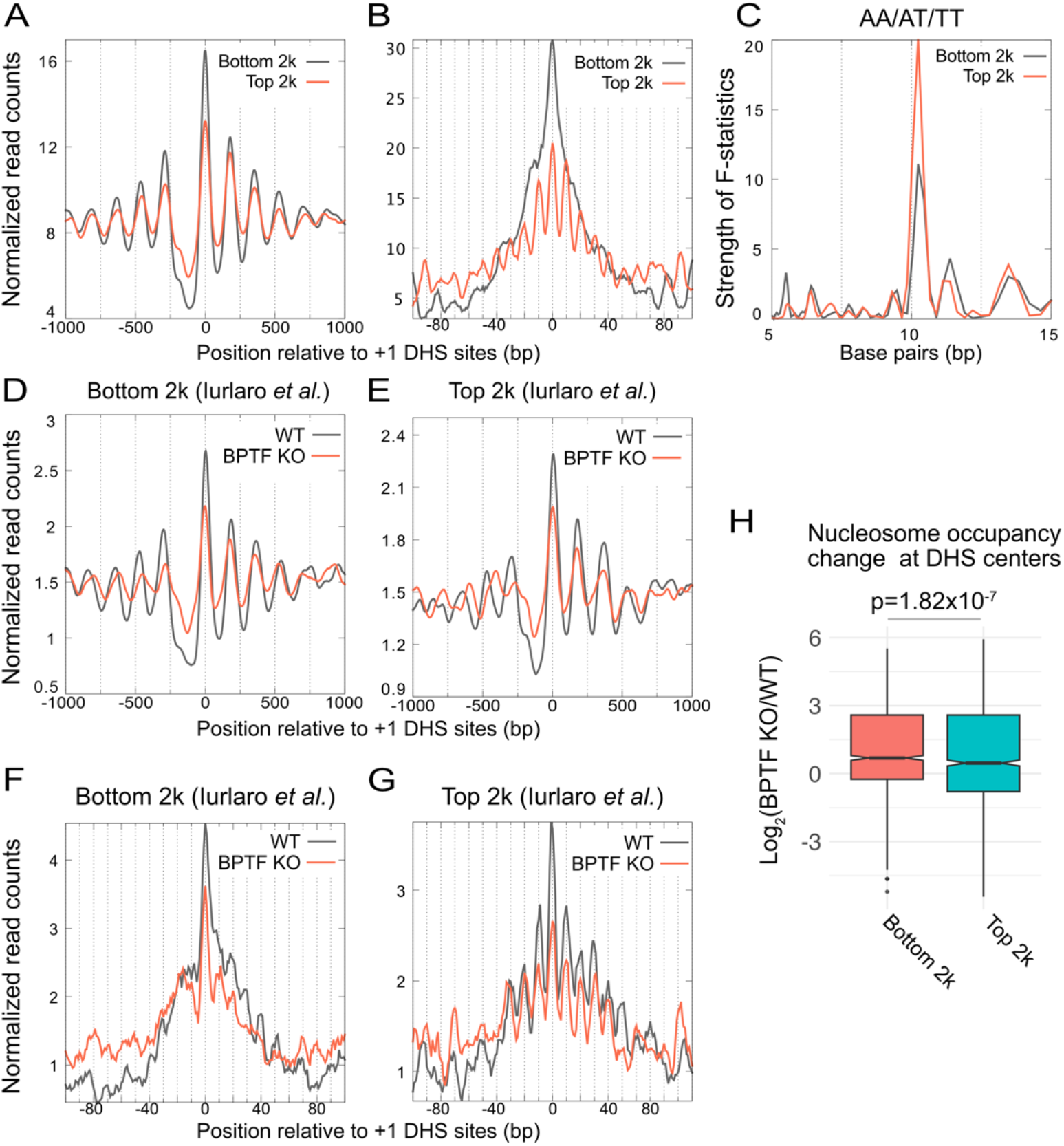
Strong rotational settings at nucleosome positions adjacent to TSS-distal DHS do not change upon loss of NURF. **A**) Low-resolution plot of mESC MNase-seq data relative to ±1000 bp of the +1 nucleosome positions (nucleosome downstream) of TSS-distal DHS. The top and bottom 2000 sites based on the strength of rotational phasing of the DHS+1 nucleosome are shown here. **B**) High-resolution plot relative to ±100 bp for the same positions as (**A**). **C**) Statistical measure of periodicity measured by multitaper of WW dinucleotide frequency for the top and bottom 2000 DHS+1 nucleosome positions shows a strong peak around 10 bp that is higher for the top compared to bottom 2000 positions. **D**) Low-resolution plot of MNase-seq data relative to ±1000 bp of the nucleosome dyad for WT and BPTF KO mESCs (from Iurlaro *et al.*) averaged over the bottom 2000 DHS+1 positions. **E**) Same as (**D**) for top 2000 DHS+1 positions. **F**) High-resolution plot of MNase-seq data relative to ±100 bp of the nucleosome dyad for WT and BPTF KO mESCs (from Iurlaro *et al.*) averaged over the bottom 2000 DHS+1 positions. **G**) Same as (**F**) for top 2000 DHS+1 positions. **H**) Log2 ratio of gain in nucleosome occupancy at TSS-distal DHS centers due to BPTF KO mESCs compared to WT mESCs represented as boxplots for the top and bottom 2000 DHS defined based on strength of rotational phasing of DHS+1 nucleosome position. P-value was calculated using Wilcoxon Rank Sum test.

With these two sets of positions, we next asked what happens to nucleosome distribution upon loss of the BPTF subunit of NURF. We analyzed published MNase-seq data generated in the presence and absence of BPTF in mESCs^9^. We observe nucleosome gain at the region upstream of DHS+1 for both weakly and strongly rotationally phased nucleosome positions upon loss of BPTF in low-res plots **(Figure 8D, E**). In high-res plots, both presence and absence of BPTF does not show any appreciable rotational phasing for the weak positions similar to our MNase-seq data presented earlier (**Figure 8F**). The alternative translational positions are clearly observable at the strong positions in WT mESC, and strikingly, these positions are preserved upon loss of BPTF (**Figure 8G**). Thus, at regions where the nucleosome landscape is regulated by NURF, rotational settings persist even after NURF is lost. We asked if strong rotational settings at DHS+1 influence the DHS upon loss of NURF. We observe significantly lower nucleosome gain at the DHS adjacent to strongly rotationally phased DHS+1 compared to weakly rotationally phased DHS+1 (**Figure 8H**). This analysis suggests that strong rotational phasing reduces invasion of the DHS site by nucleosomes upon BPTF loss. In summary, mESCs similar to budding yeast have a significant fraction of nucleosome positions with strong rotational settings, and the rotational settings correlate with strength of WW oscillations at 10 bp. Further, similar to budding yeast, loss of NURF results in quantized 10 bp shift in nucleosome positions.

## DISCUSSION

Here, we developed computational methods to quantify the strength of rotational setting at individual nucleosome positions, and to unlock bp-resolution positioning information even from lower-resolution methods like MNase-seq. This allowed us to show that sequence-directed rotational positioning, though not directing translational positions, maintains the solvent-accessible face of the DNA helix even when nucleosomes are repositioned by remodeling and transcription elongation. We find that nucleosome positions with stronger rotational settings have stronger A/T and G/C dinucleotide periodicity every 10 bp, agreeing with long-established rules for nucleosome positioning, and that strong rotational settings lend themselves to sampling of alternate translational positions, possibly due to the equivalence of sequence rules at every strong histone-DNA contact. This was consistent with our observation that *in vitro* SGD nucleosomes had the same rotational settings as those *in vivo*, and, based on the strength of 10 bp WW frequency, that strong rotational settings appear even in sequences derived from species without histones when introduced into yeast. These observations support the model that the sequence sets up a shallow energy potential that influences nucleosome mobility *in vitro* and *in vivo*^51^. Although this energy-landscape model has long been proposed, our systematic perturbation analysis provides a direct *in vivo* test of it: across TKO, loss of INO80 and RSC, inhibition of transcription elongation, and in mammalian cells, the rotational setting is preserved while translational preferences shift, as the model predicts.

Chromatin regulators therefore appear to preserve, rather than alter, the face of DNA contacting histones, and nucleosome movements due to different classes of regulators (different remodeler families, elongation factors, and RNA polymerase) are all quantized to an apparent 10 bp step. We propose that the sequence-encoded rotational minima act as a molecular detent: like a mechanical detent that permits motion but clicks into evenly spaced stops, they allow nucleosomes to be moved in either direction while favoring stable positions every ∼10 bp that preserve the rotational setting.

Our observation of ∼10 bp quantization in steady-state nucleosome positions may appear to conflict with structural studies showing that remodelers translocate DNA in 1-2 bp steps^52,53^. However, these observations are reconcilable: remodelers likely move nucleosomes continuously along DNA, but the sequence-encoded rotational energy landscape creates energetic minima every ∼10 bp. When remodeler activity ceases (due to dissociation, ATP depletion, or competition with other factors), nucleosomes relax to the nearest favorable rotational position^54^. Thus, the detent mechanism describes the preferred stopping points rather than the step size of translocation itself. This framework explains why population-level mapping consistently reveals 10 bp periodicity while single-molecule studies observe continuous movement.

Nucleosomes have been long known to have intrinsic sequence preferences, but chromatin regulators *in vivo* are thought to override these intrinsic sequence preferences. Our analysis here shows that the intrinsic preferences mainly manifest as strong rotational settings at most nucleosome positions genome-wide. These rotational settings are not overridden by chromatin regulators but seem to be co-opted by them when they move nucleosomes. At the preferred rotational setting, a flatter preference between alternative translational positions *in vitro* is sharpened *in vivo*, with this sharpening highest at +1 nucleosome and weakening with increasing distance downstream from the TSS. This agrees well with statistical positioning driving the preferred translational positions, with promoter having presumably strong barriers, and remodelers when pushing against this barrier select preferred translational positions. The promoter barrier, likely established by sequence-directed nucleosome depletion and transcription factor binding, provides a fixed reference point from which remodelers position nucleosomes. Critically, this barrier can only select among the translational positions that preserve intrinsic rotational settings; it cannot override sequence preferences. This explains why +1 nucleosomes show the strongest positioning: they experience the combined influence of a nearby barrier and concentrated remodeler activity, selecting the single rotational-compatible position closest to the TSS^21^. The gradual attenuation of positioning with distance from the TSS reflects both weaker barrier influence and increased competition among multiple rotational-compatible positions.

The universality of rotational constraints across remodelers, transcription, and organisms suggests evolutionary pressure to maintain this regulatory layer. The positioning of transcription factor binding motifs relative to rotational settings may be under selection, as the face of DNA exposed to solution determines accessibility to regulatory proteins^20,33^. Additionally, rotational positioning could facilitate nucleosome-mediated cooperativity between distantly bound factors by ensuring they occupy the same helical face. These considerations suggest that apparent ‘fuzziness’ in nucleosome positioning at low resolution may actually represent functional sampling among discrete, rotationally equivalent positions.

Budding yeast has a higher A/T content, and we wondered if the sequence composition of the genome might be biasing our results given the known A/T bias of MNase. We rule out MNase bias or genome sequence content giving rise to the patterns we interpret as alternative translational positions with the same rotational settings by two analyses. First, digestion of naked yeast genome does not give rise to observed patterns, ruling out MNase bias. Second, the mouse genome has a lower A/T content but still shows clear preference for a specific rotational setting with the same preference for dinucleotide frequencies. Similar to yeast, loss of remodeler function leads to nucleosome gain at DHS sites but not change in rotational settings. Finally, even sequences from species without histones, when introduced into yeast, display rotational settings commensurate with the strength of their 10 bp oscillations of WW frequency, with no clear correlation to A/T content. Thus, our model of shallow nucleosome binding potential that dictates rotational settings might be applicable widely. Interestingly, the nucleosome landscape in human cells has also been shown to prefer 10 bp periodic patterns in dinucleotide frequencies^55,56^. Remodelers can move nucleosomes in 1 bp steps following multi-bp steps^52,53^; it would be critical for future studies to characterize how remodeler steps interface with preferred rotational positions. Future questions could also involve asking whether preferred rotational settings are overridden during processes like replication-coupled nucleosome deposition or identifying species where our model is not applicable. These exceptions would identify chromatin regulators that do override intrinsic sequence preferences of nucleosomes and the processes that require such activity.

## ACKNOWLEDGMENTS

We thank Dr. Neelanjan Mukherjee for suggesting the idea to apply multitaper to the datasets. We thank Dr. Aaron Johnson, Dr. Catherine Musselman, Dr. Jay Hesselberth, Dr. Sujatha Jagannathan, and members of the Ramachandran lab for critical reading of the manuscript. We would like to acknowledge support from NIH grant R35GM156411 (S. R.), the Pew Charitable Trusts and the Alexander and Margaret Stewart Trust (S.R.).

## METHODS

### Data Availability

MNase-seq data generated in mESCs for this study are available at GEO (GSE314045).

### Mouse Embryonic Stem Cell Culture

Mouse embryonic stem cell line E3 (mESCs; Gates Biomanufacturing Facility at University of Colorado Denver) were utilized for MNase-seq experiments. Cells were tested for mycoplasma contamination and confirmed to be negative before experimental work. mESCs were cultured under feeder-free conditions on 0.1% gelatin-coated T-25 flasks in knockout DMEM (*Life Technologies*), 20% embryonic stem cell-qualified fetal bovine serum (FBS; *Life Technologies*), 2 mM L-glutamine (*Life Technologies*), 0.1 mM non-essential amino acids (*Life Technologies*), 100UmL^-1^ penicllin/streptomycin (*Life Technologies*), 0.05 mM β-mercaptoethanol (*Sigma*), and 1000UmL^-1^ ESGRO mouse leukemia inhibitory factor medium supplement (LIF; *Millipore*). mESCs were cultured at 37°C with 5% CO_2_ and subcultured every 2 days. Cells were harvested for experiments by aspirating the growth media, applying 0.25% trypsin-EDTA (Gibco) for 2-3 minutes, and centrifuging at 2000 rpm for 5 minutes.

### MNase-seq for Mouse Embryonic Stem Cells

After harvesting serum-grown mESCs, cells were washed in cold 1x phosphate buffered saline solution (PBS). Cell number and viability were measured with Countess^TM^ 3 Automated Cell Counter (*Invitrogen*). MNase-seq digestions proceeded with 1 x 10^6^ mESCs. mESC samples were centrifuged at 2000 rpm for 5 minutes to remove 1x PBS. Cells were resuspended in 1 mL cold Nuclear Extraction Buffer (20 mM HEPES pH 8.0, 10 mM KCl, 0.1% Triton, 20% glycerol, 0.5 mM spermidine) and incubated on ice for 10 minutes to facilitate nuclei isolation. Nuclei were centrifuged at 2500 rpm for 10 min, then resuspended in 40 uL Reaction Buffer (20 mM HEPES pH 7.5, 150 mM NaCl, 0.5 mM spermidine, 0.01% digitonin, 2 mM EGTA, 10 mM CaCl_2_, 10 mM MgCl_2_). Samples were incubated at 37°C for 5 minutes prior to starting digestion to allow for reaction temperature equilibration. 0.07 U micrococcal nuclease (*Sigma*) was added to each sample along with nuclease-free water (*Invitrogen*) to adjust reaction volumes to 50 uL. Reactions proceeded at 37°C for time points of 3, 6, 9, 12, and 15 minutes. At each time point, 10 uL of reaction (0.2 x 10^6^ mESC nuclei) was removed and quenched with 10 uL 2x Stop Buffer (500 mM NaCl, 10 mM EGTA, 0.1% digitonin, 1% SDS, 100 ug RNase A, 50 ug glycogen). Digested DNA was purified by phenol:chloroform:isoamyl alcohol extraction protocol. Purifed DNA concentration and sample quality was determined via Qubit 4 fluorometer and Agilent D1000 Tapestation respectively.

Sequencing libraries were prepared using the SRSLY PicoPlus NGS Library Prep Kit and protocol, described as follows. 10 ng of digested DNA was combined with SRSLY ssEnhancer and incubated on ice for 2 minutes, followed by heat shock at 98°C for 3 minutes and another incubation on ice for 2 minutes. Phosphorylation and ligation reaction was achieved by mixing sample with SRSLY Adapter A, SRSLY Adapter B, and SRSLY Master Mix and then incubating samples at 37°C for 1 hour. Samples were then purified in a final volume of 15 uL nuclease-free water via 0.54x AMPure bead clean to enrich for small fragments. To each sample, SRSLY PCR Master Mix and unique P5 and P7 barcoded primers were added. The following PCR protocol was used: 98C for 3 minutes, 98°C for 20 seconds, 65°C for 30 seconds, 72°C for 30 seconds, 72°C for 1 minute with steps 2-4 repeated for 10 cycles. Final library purification was achieved via 0.95x AMPure bead clean, after which library quality was assessed via Agilent TapeStation.

### Data analysis

#### Yeast datasets

Published datasets (H3-Q85C cleavage mapping, MNase-seq from WT, TKO, SGD, SGD+Remodelers, naked DNA, RNA Pol II anchor-away, INO80 depletion, Sth1 anchor-away, Spt4 deletion, BPTF knockout, and yeast artificial chromosomes (YACs) hybrids) were obtained from Gene Expression Omnibus database (GEO, **Supplementary Table 1**). For yeast datasets, the paired-end fastq files were mapped to the *S. cerevisiae* genome (sacCer3) using Bowtie2^57^. Duplicate reads with the same start and end coordinates were not included in the downstream analysis and discarded. Bam files were created using samtools^58^, followed by bedpe generation using sam2bam command in bedtools^59^. From the bedpe file, the extreme ends of the mates representing the ends of the sequenced fragments were extracted into a bed file, where each entry contained the chromosome, start and end coordinate of each unique sequenced fragment. Centers of fragments belonging to different length classes (140-160 bp for low-res and 142-152 bp for high-res, 48-52 bp for H3-85C dataset) were used to make coverage files in the wiggle format with a step size of 1 bp. Nucleosome positions^25^ relative to the transcription start sites^60^ were defined following previous studies^61^. The coverage of fragment centers at each base-pair location was normalized by the factor N:

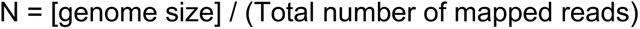

Here, [genome size] refers to a number close to total mappable bases for each strain (sacCer3 = 12157105, Yeast + Mmyco = 1222180, Yeast + Mpneumo = 817920, Yeast + Vitis = 277240, Yeast + Falciparum = 91204).

#### SGD Datasets

SGD experiments in the original study were performed on a subset of plasmids covering the yeast genome. To identify genomic regions covered by the plasmids used in the study, we performed domain calling adapted from Ref.^62^ on the SGD+Chd1 dataset and analyzed only those nucleosome positions that overlap with regions with coverage.

#### Calling nucleosome peaks in yeast hybrid strains

For the bacterial and parasitic genome sequences that were added as an extra chromosome in the yeast hybrid strains, chemical cleavage mapping data does not exist. To overcome this limitation, we developed a high-resolution nucleosome calling algorithm. We first called peaks across the genome using our peak caller (https://github.com/satyanarayan-rao/tf_nucleosome_dynamics/blob/main/CUTnRUN_Peak_Calling/call_peaks_iterative.py) for each strain using the wig file generated from 142-152 bp fragments as the input. We then scanned 10 bp upstream and downstream of the peaks in the high-res nucleosome coverage map to identify local maxima and adjusted the original nucleosome position to obtain a high-resolution nucleosome position.

#### Low-res and high-res plots

For MNase-seq data, we generated two versions of genome-wide coverage files in the wiggle format, 140-160 bp for low-res and 142-152 bp for high-res. Low-res plots were generated using coverage of fragment centers of 140-160 bp fragments with a smoothing window of 10 bp. High-res plots were generated using coverage of fragment centers of 142-152 bp with a smoothing window of 2 bp.

#### Mouse datasets

Nucleosome center positioning score (NCP score) in mm9 coordinates from Voong *et al*.^26^ were remapped to mm10 coordinates using liftOver (UCSC Genome Browser)^63^. Mouse ENCODE DHS dataset (ENCFF048DWN) was used to identify TSS-distal DHSs^64,65^.

For published and inhouse mouse MNase-seq datasets, the paired-end fastq files were mapped to the *M. musculus* genome (mm10) using Bowtie2^57^. Duplicate reads with the same start and end coordinates were not included in the downstream analysis and discarded. Bam files were created using samtools^58^, followed by bedpe generation using sam2bam command in bedtools^59^. From the bedpe file, the extreme ends of the mates representing the ends of the sequenced fragments were extracted into a bed file, where each entry contained the chromosome, start and end coordinate of each unique sequenced fragment. Centers of fragments belonging to different length classes (140-160 bp for low-res and 142-152 bp for high-res, 48-52 bp for H3-85C dataset) were used to make coverage files in the wiggle format with a step size of 1 bp. The coverage of fragment centers at each base-pair location was normalized by the factor N:

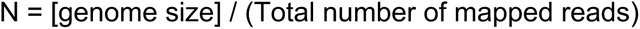

Here, [genome size] refers to a number close to total mappable bases, which was 2.8×10^9^ for mouse.

#### Multitaper Analysis of rotational phasing

Coverage of nucleosome centers (from 147±5 fragments from MNase-seq or 50±2 bp fragments from H3-Q85C chemical cleavage mapping) at a 100 bp window around a given nucleosome position was used as a “time series” input for the multitaper package in R^29,66^.

#### Multitaper command parameters

Multitaper outputs an F-statistic as a function of frequency. The maximum F-statistic in the window of frequency that would roughly correspond to 8-13.5 bp was extracted as the strength of the rotational setting for that nucleosome position. Following methodology presented by Calviello *et al.*^29^, we calculated P values from the *F*-statistic^67^ by using 2 and 2*k* – 2 degrees of freedom where *k* is the number of base pairs (10 bps in this study), giving a cut-off of 3.2 for a F-statistic with p≈0.05^29^.

#### Multitaper Analysis for dinucleotide frequency

The sequences for groups of nucleosome positions were aligned based on dyad locations and dinucleotide frequencies (either AA/AT/TA/TT (“WW”) or GG/GC/CG/CC (“SS”)) were calculated +/-73 bp from the dyad. This WW or SS dinucleotide frequency was used as a “time series” input for the multitaper package in R. The F-statistic as a function of frequency^-1^ (so in units of bp) was plotted for different nucleosome sets.

#### Fractional Enrichment at Rotational Positions and Dinucleotide Enrichment at Rotational Positions

Per-nucleosome occupancy and sequence preference were quantified using two complementary scores, Fractional Enrichment at Rotational Positions (FERP) and Dinucleotide Enrichment at Rotational Positions (DERP). Both scores were defined relative to the same set of empirically detected rotational peak and trough positions (**Supplementary Figure S1B, C**), ensuring direct comparisons between the nucleosome density and the underlying sequence signal at every nucleosome locus. Nucleosome and dinucleotide peak and trough positions were determined from the population-mean coverage and dinucleotide profiles of the top 20% of nucleosomes ranked by H3-Q85C F-statistics (the reference subset). Peak detection was performed using scipy.signal.find_peaks^68^ with a minimum inter-peak distance of 9 bp and a prominence threshold of 0.01, followed by an iterative gap-filling procedure to recover closely spaced peak pairs. Trough positions were assigned as the minimum of the population-mean profile between each adjacent pair of detected peaks. Once determined from the H3-Q85C Chemical Cleavage reference dataset, these positions were saved and applied identically to all other experimental conditions to ensure cross-condition comparability. For each detected peak, the mean coverage over a 5 bp window centered on the peak position (center mean) and the mean of the 5 bp windows at the nearest flanking trough positions on either side (flank mean) are calculated. The FERP score for a given nucleosome is defined as the mean of the center-minus-flank differences across all detected peak positions. Nucleosomes with fewer than five non-zero peak positions were excluded. For calculating the DERP score, each base-pair position in the nucleosome window is assigned a dinucleotide score of +1 (WW: AA, TT, AT, TA), -1 (SS: GG, CC, GC, CG), or 0 (mixed). The DERP score is computed as the mean 5bp window score at detected peak positions minus the 5bp window score at detected trough positions.

## Declaration of generative AI and AI-assisted technologies in the manuscript preparation process

During the preparation of this work the authors used Claude.ai to improve language and readability and to assist with the development of analysis code for the implementation of the FERP and DERP scoring routines. After using this tool, the authors reviewed, tested, and edited the content and code as needed and take full responsibility for the content of the published article.

