## Supplemental Figures for "Rotational settings quantize nucleosome movement by chromatin regulators"

**Supplementary Table 1**. GEO Accession numbers for all published datasets reanalyzed in each figure.

| Figures | GEO Accession Number |
|---|---|
| Figure 1 | GSE97290, GSE59523 |
| Figure 2 | GSE209681 |
| Figure 3 | GSE209681 |
| Figure 4 | GSE217022 |
| Figure 5 | GSE141007, GSE118214 |
| Figure 6 | GSE141007, GSE159291 |
| Figure 8 | GSE234251 |
| Figure S1 | GSE97290, GSE59523 |
| Figure S2 | GSE97290, GSE216450, GSE59523 |
| Figure S3 | GSE216450, GSE209681 |
| Figure S4 | GSE141007 |
| Figure S5 | GSE234251, GSE118214 |
| Figure S6 | GSE234251, GSE118214 |
| Figure S7 | GSE234251, GSE159291 |

### Figure S1

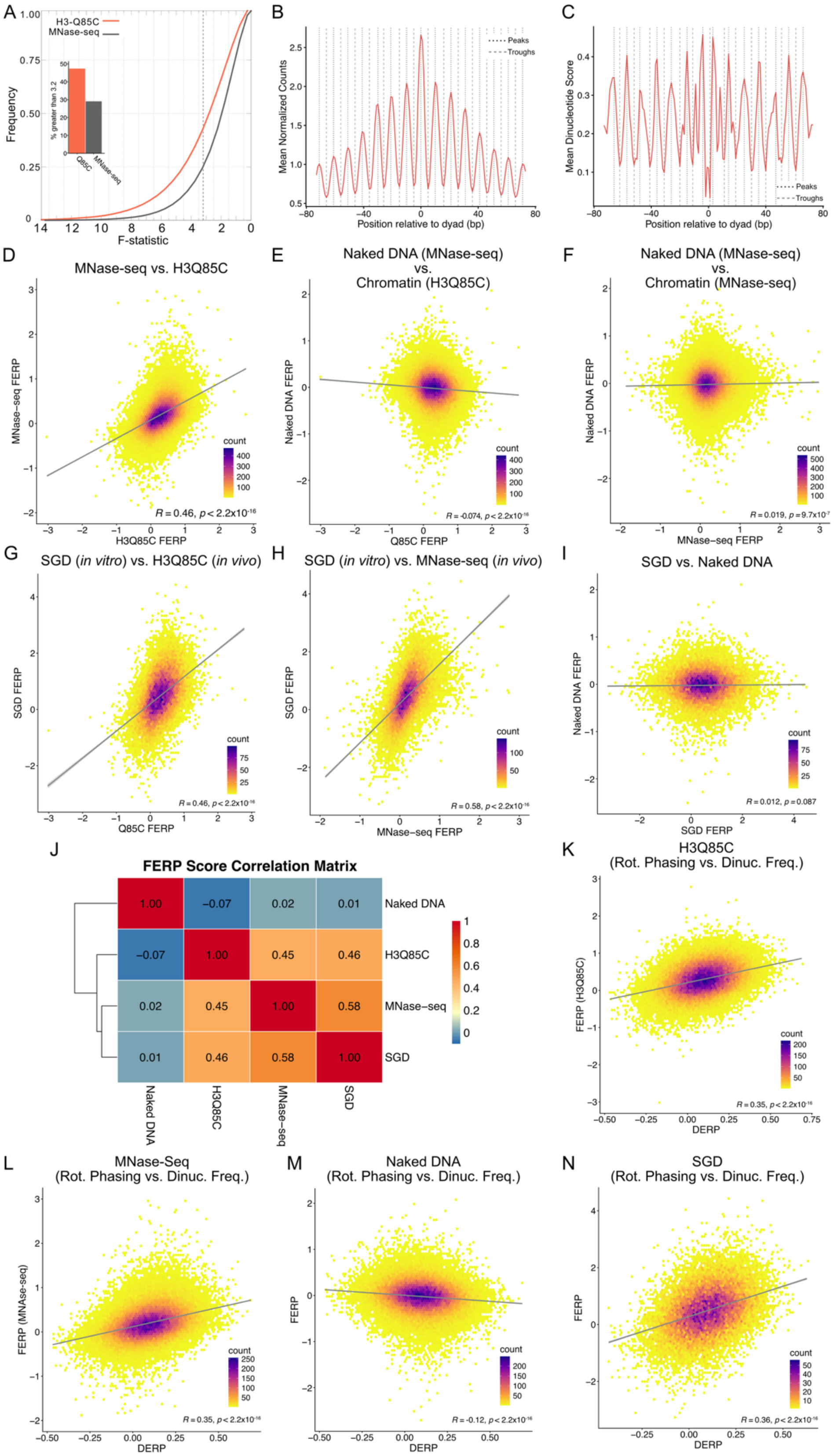
A
H3-Q85C
MNase-seq
Frequency
F-statistic
% greater than 3.2
Q85C
MNase-seq
B
Mean Normalized Counts
Position relative to dyad (bp)
Peaks
Troughs
C
Mean Dinucleotide Score
Position relative to dyad (bp)
Peaks
Troughs
D
MNase-seq vs. H3Q85C
MNase-seq FERP
H3Q85C FERP
count
R = 0.46, p < 2.2x10-16
E
Naked DNA (MNase-seq) vs. Chromatin (H3Q85C)
Naked DNA FERP
Q85C FERP
count
R = -0.074, p < 2.2x10-16
F
Naked DNA (MNase-seq) vs. Chromatin (MNase-seq)
Naked DNA FERP
MNase-seq FERP
count
R = 0.019, p = 9.7x10-7
G
SGD (in vitro) vs. H3Q85C (in vivo)
SGD FERP
Q85C FERP
count
R = 0.46, p < 2.2x10-16
H
SGD (in vitro) vs. MNase-seq (in vivo)
SGD FERP
MNase-seq FERP
count
R = 0.58, p < 2.2x10-16
I
SGD vs. Naked DNA
Naked DNA FERP
SGD FERP
count
R = 0.012, p = 0.087
J
FERP Score Correlation Matrix
Naked DNA
H3Q85C
MNase-seq
SGD
K
H3Q85C
(Rot. Phasing vs. Dinuc. Freq.)
FERP (H3Q85C)
DERP
count
R = 0.35, p < 2.2x10-16
L
MNase-Seq
(Rot. Phasing vs. Dinuc. Freq.)
FERP (MNase-seq)
DERP
count
R = 0.35, p < 2.2x10-16
M
Naked DNA
(Rot. Phasing vs. Dinuc. Freq.)
FERP
DERP
count
R = -0.12, p < 2.2x10-16
N
SGD
(Rot. Phasing vs. Dinuc. Freq.)
FERP
DERP
count
R = 0.36, p < 2.2x10-16

**Figure S1. Large fraction of yeast nucleosomes is rotationally phased. A**) Cumulative frequency of fraction of nucleosome positions as a function of decreasing F-statistic from multitaper for all nucleosome positions in the yeast genome. The inset shows a bar-plot with % of nucleosome positions with F-statistic > 3.2. **B**) Average H3Q85C chemical cleavage mapping data (50±2 bp) for the top quintile of rotational positioning, plotted relative to dyad position. The dotted lines indicate peaks and dashed lines indicate troughs. **C**) Average dinucleotide score (setting WW=1, SS=-1, WS/SW=0) for the top quintile of rotational positioning, plotted relative to dyad position. The dotted lines indicate peaks and dashed lines indicate troughs. **D-I**) Hexagonal binning comparing FERP scores for specified datasets. Pearson correlation coefficients are shown for each comparison. **J**) FERP score correlation matrix comparing Naked DNA, H3Q85C, MNase-seq and SGD. **K-N**) Hexagonal binning comparing FERP scores (rotational phasing) for specified datasets versus Dinucleotide Enrichment at Rotational Positions (DERP) scores (dinucleotide frequencies). Pearson correlation coefficients are also shown for each comparison.

**Figure S2**

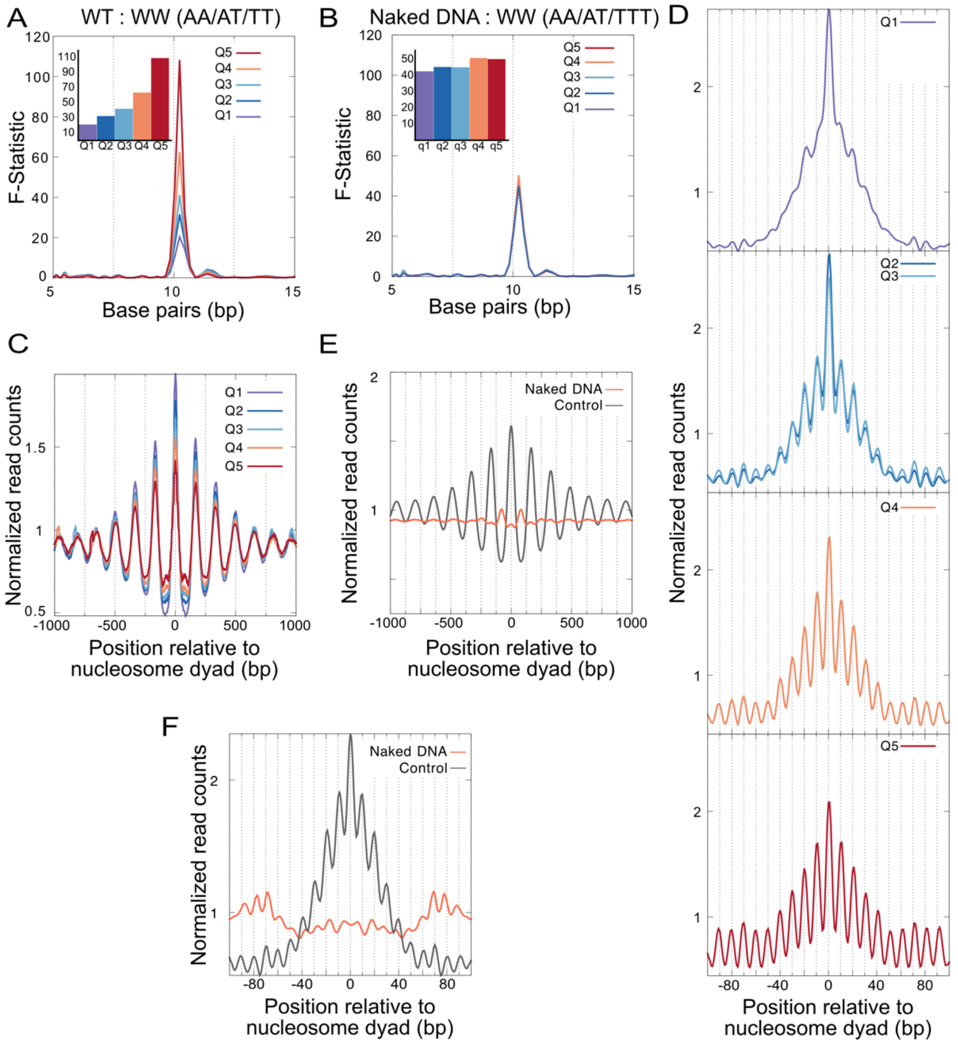


**Figure S2. Yeast rotational positions do not arise from MNase bias. A**) Statistical measure of periodicity measured by multitaper of WW dinucleotide frequency for nucleosome positions in each quintile from MNase-seq shows a strong peak around 10 bp that is higher for higher quintiles. Inset shows a bar plot for the value of the F-statistic at the peak around 10 bp. **B**) Same plot as (**A**) for dataset generated after MNase digestion of naked DNA does not show a trend of stronger 10-bp periodicity of WW dinucleotides with higher quintiles. **C**) Low-resolution plot of normalized read counts from MNase-seq data averaged across nucleosome positions divided into quintiles based on the strength of rotational setting (F-statistic from multitaper), plotted ±1000 bp relative to nucleosome center. (**D**) High-resolution plot of normalized read counts from MNase-seq data averaged across nucleosome positions divided into same quintiles as (**C**). **E**) Low-resolution plot of normalized read counts from MNase-seq data averaged across all nucleosome positions genome-wide compared to similar plot from the dataset generated by MNase digestion of naked DNA, plotted 1000 bp relative to nucleosome center. **F**) High-resolution plot of normalized read counts from MNase-seq data averaged across all nucleosome positions genome-wide compared to similar plot from the dataset generated by MNase digestion of naked DNA, plotted ±100 bp relative to the nucleosome center.

**Figure S3**

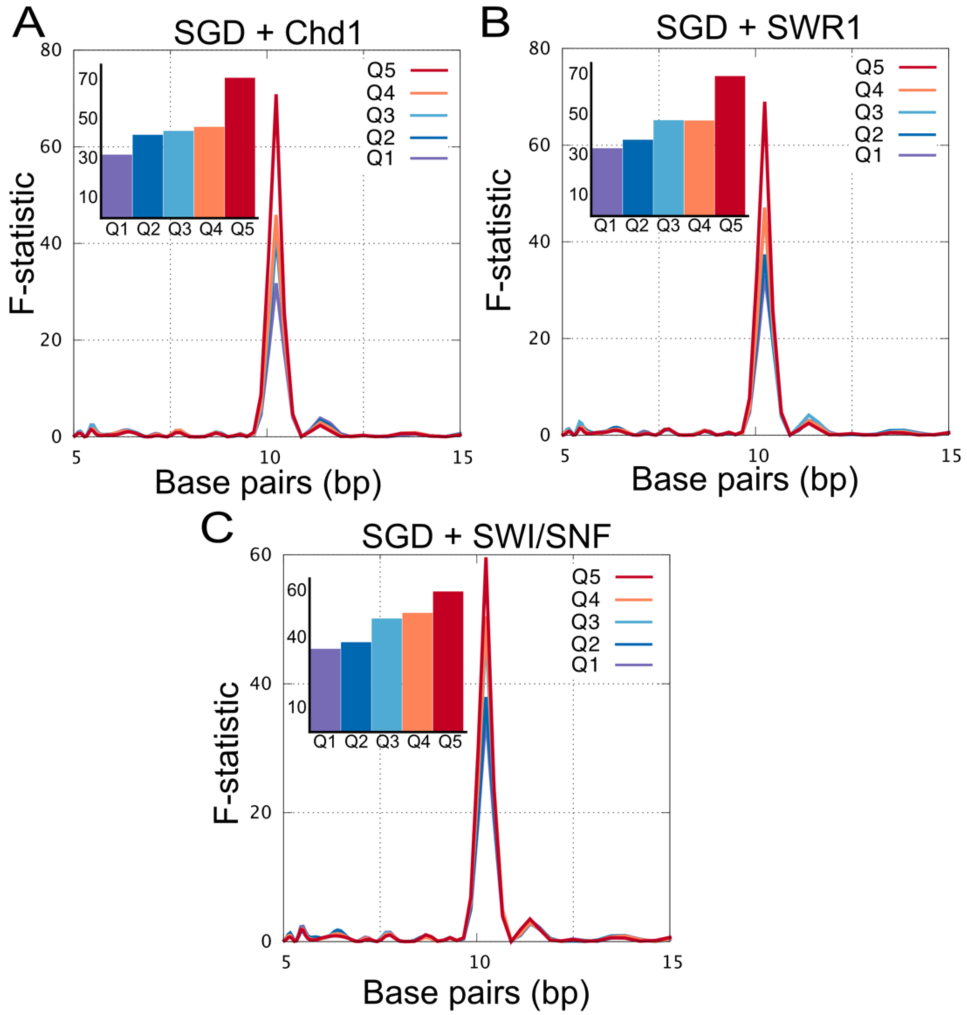


**Figure S3. Intrinsically preferred rotational settings preserved by remodelers *in vitro.*** (**A-C**) Statistical measure of periodicity measured by multitaper of WW dinucleotide frequency for nucleosome positions in each quintile shows a strong peak around 10 bp that is higher for higher quintiles. Inset shows a bar plot for the value of the F-statistic at the peak around 10 bp.

#### Figure S4

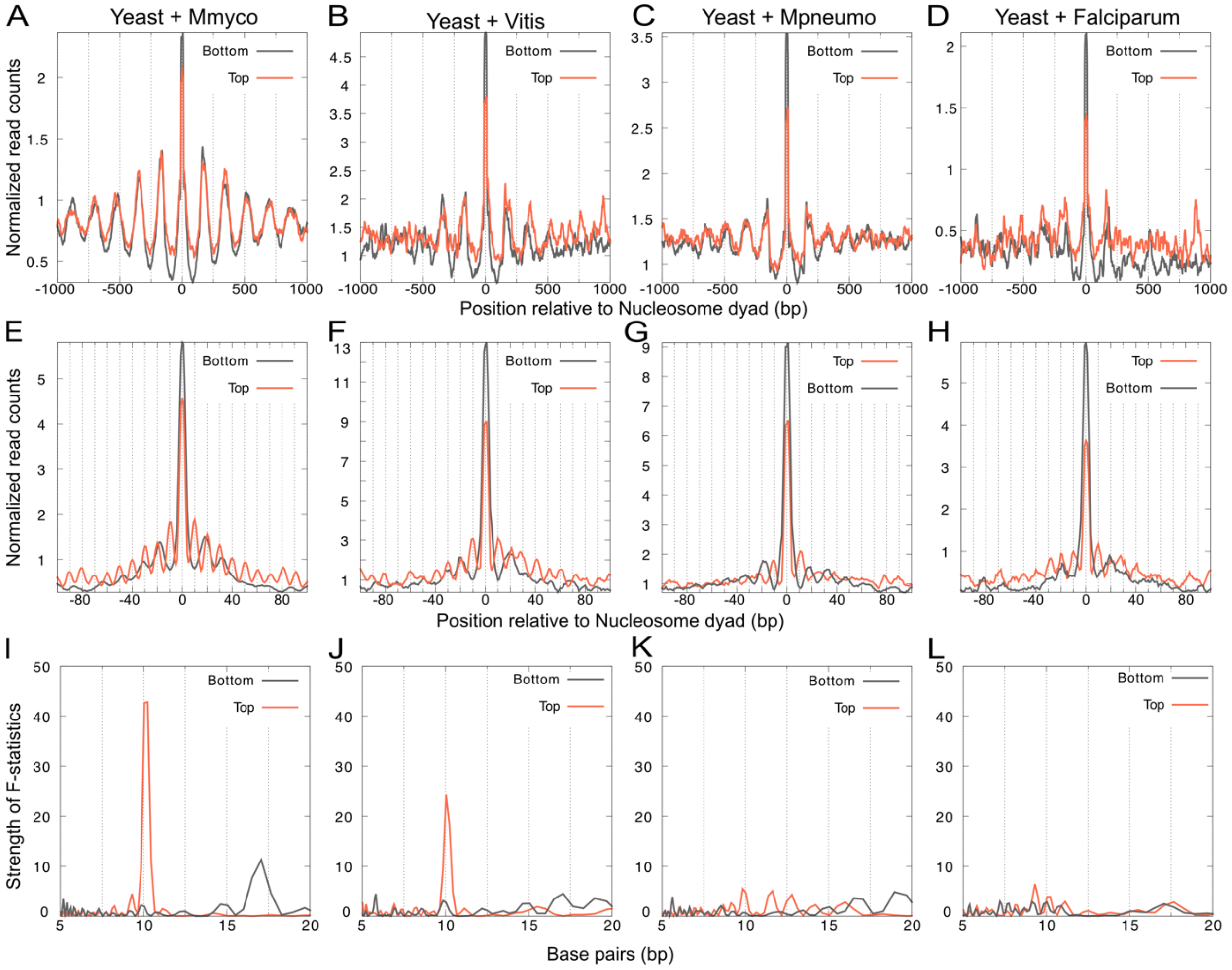


**Figure S4. Strength of rotational positioning correlates with the strength of WW periodicity in foreign DNA inserted into yeast genome.** (**A-D**) Low-resolution plot of MNase-seq data derived from yeast hybrid strains strain combinations: Yeast + Mmyco (DNA from *Mycoplasma mycoides*), Yeast + Vitis (DNA from *Candidatus Phytoplasma vitis*), Yeast + Mpneumo (DNA from *Mycoplasma pneumoniae*), and Yeast + Falciparum (DNA from *Plasmodium falciparum*), at top and bottom 10% of nucleosome positions based on strength of rotational positioning, identified on the exogenous DNA sequences. (**E-H**) High-resolution plot of plot of MNase-seq data for same strains and nucleosome positions as (**A-D**). (**I-L**) Statistical measure of periodicity measured by multitaper of WW dinucleotide frequency for top and bottom 10% of nucleosome positions based on strength of rotational positioning shows a strong peak around 10 bp for the top 10% of positions of Mmyco and Vitis, which also show strong rotational positioning.

**Figure S5**

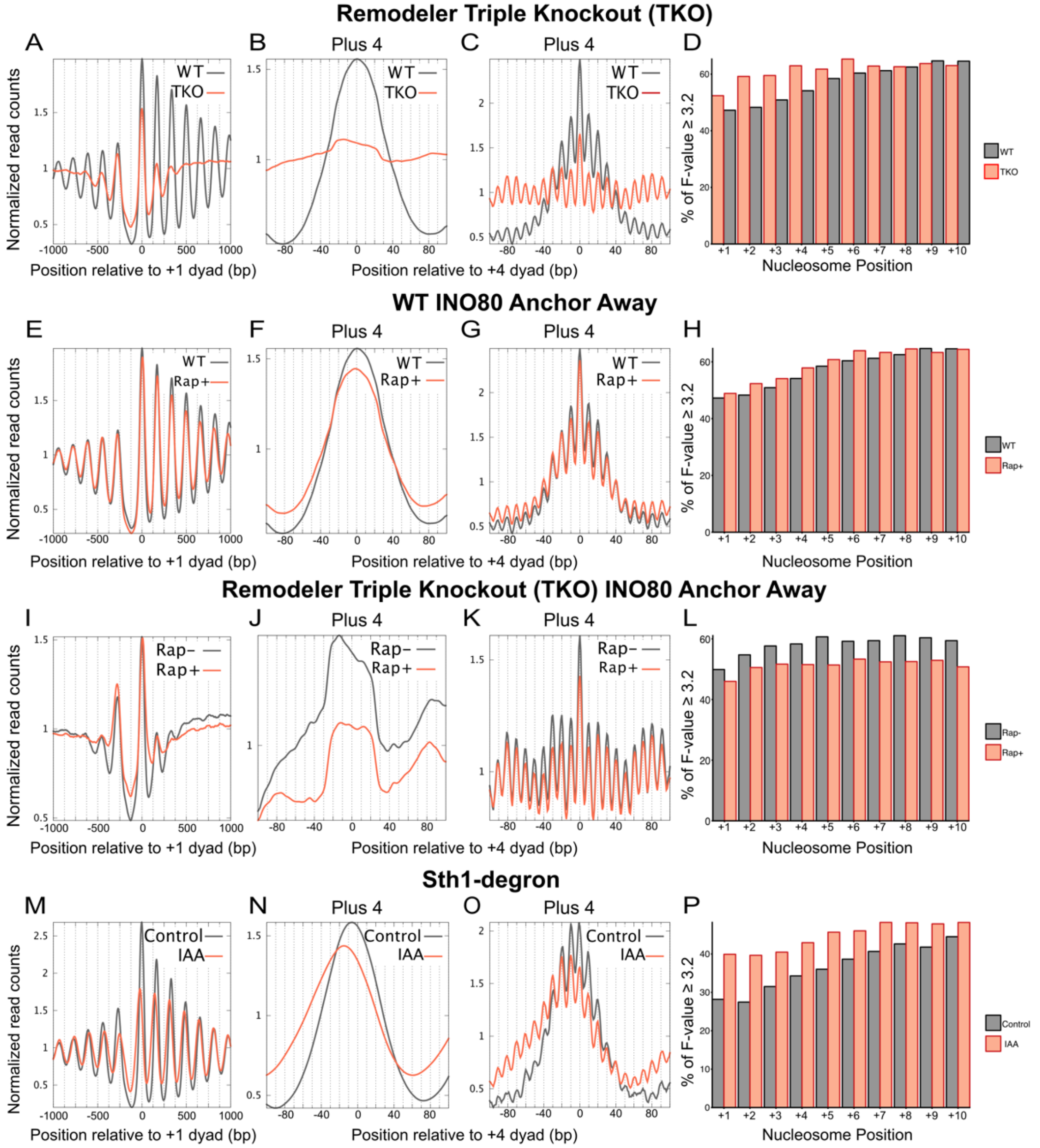


**Figure S5. Nucleosome shifts due to perturbations in remodeling are quantized.** (**A**) Low-resolution plot of MNase-seq data derived from TKO strain compared to WT at +/-1000 bp relative to the +1 nucleosome position across all genes. (**B**) Low-resolution plot of MNase-seq data derived from TKO strain compared to WT at +/-100 bp relative to the +4 nucleosome position across all genes. (**C**) High-resolution plot of MNase-seq data derived from TKO strain compared to WT at +/-100 bp relative to the +4 nucleosome position across all genes. (**D**) Fraction of genes with significant rotational phasing (F-statistic > 3.2) at each nucleosome position downstream of the TSS based on the MNase-seq data derived from TKO strain compared to WT. (**E-H**) Same as (**A-D**) comparing INO80 anchor-away strain ±rapamycin treatment for 90 minutes. (**I-L**) Same as (**A-D**) for INO80 anchor-away strain in the TKO background with 90-minute rapamycin treatment compared to no rapamycin. (**M-P**) Same as (**A-D**) comparing Sth1-degron strain ±IAA treatment for 60 minutes.

**Figure S6**

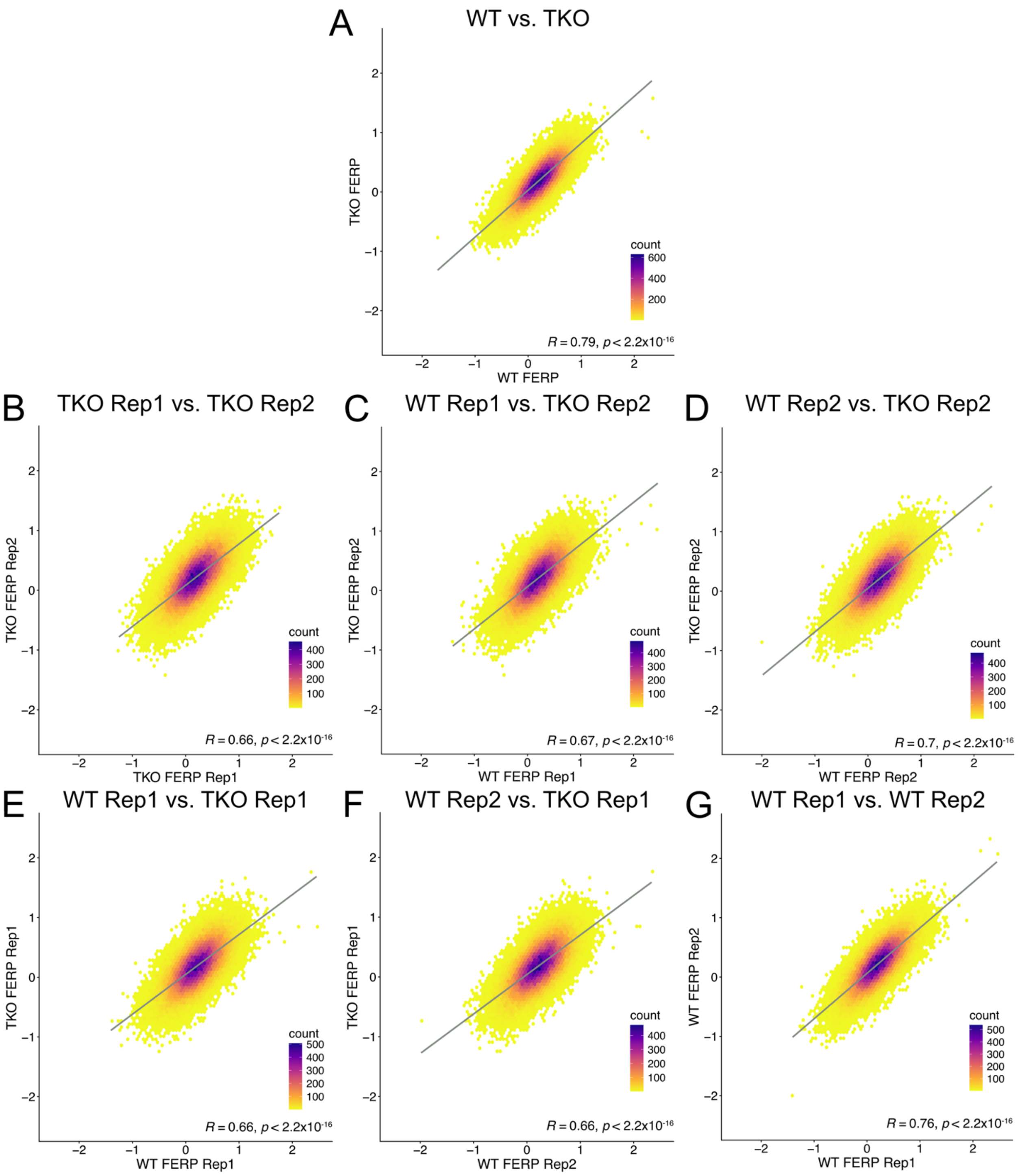


**Figure S6. Rotational positioning is highly similar between WT and TKO. A**) Hexagonal binning of combined replicate FERP scores for WT versus TKO strains shows highly significant correlation (r=0.79). **B**) Hexagonal binning of FERP scores to determine within-condition reproducibility of TKO across replicates (Replicate 1 versus Replicate 2, r=0.66). **C**) Hexagonal binning of FERP scores for cross-condition comparison of WT Replicate 1 versus TKO FERP

Replicate 2 (r=0.67). **D**) Hexagonal binning of FERP scores for cross-condition comparison of WT Replicate 2 versus TKO Replicate 2 (r=0.7). **E**) Hexagonal binning of FERP scores for cross-condition comparison of WT Replicate 1 versus TKO Replicate 1 (r=0.66). **F**) Hexagonal binning of FERP scores for cross-condition comparison of WT Replicate 2 versus TKO Replicate 1 (r=0.66). **G**) Hexagonal binning of FERP scores for within-condition reproducibility of WT scores biological replicates (Replicate 1 versus Replicate 2, r=0.76).

**Figure S7**

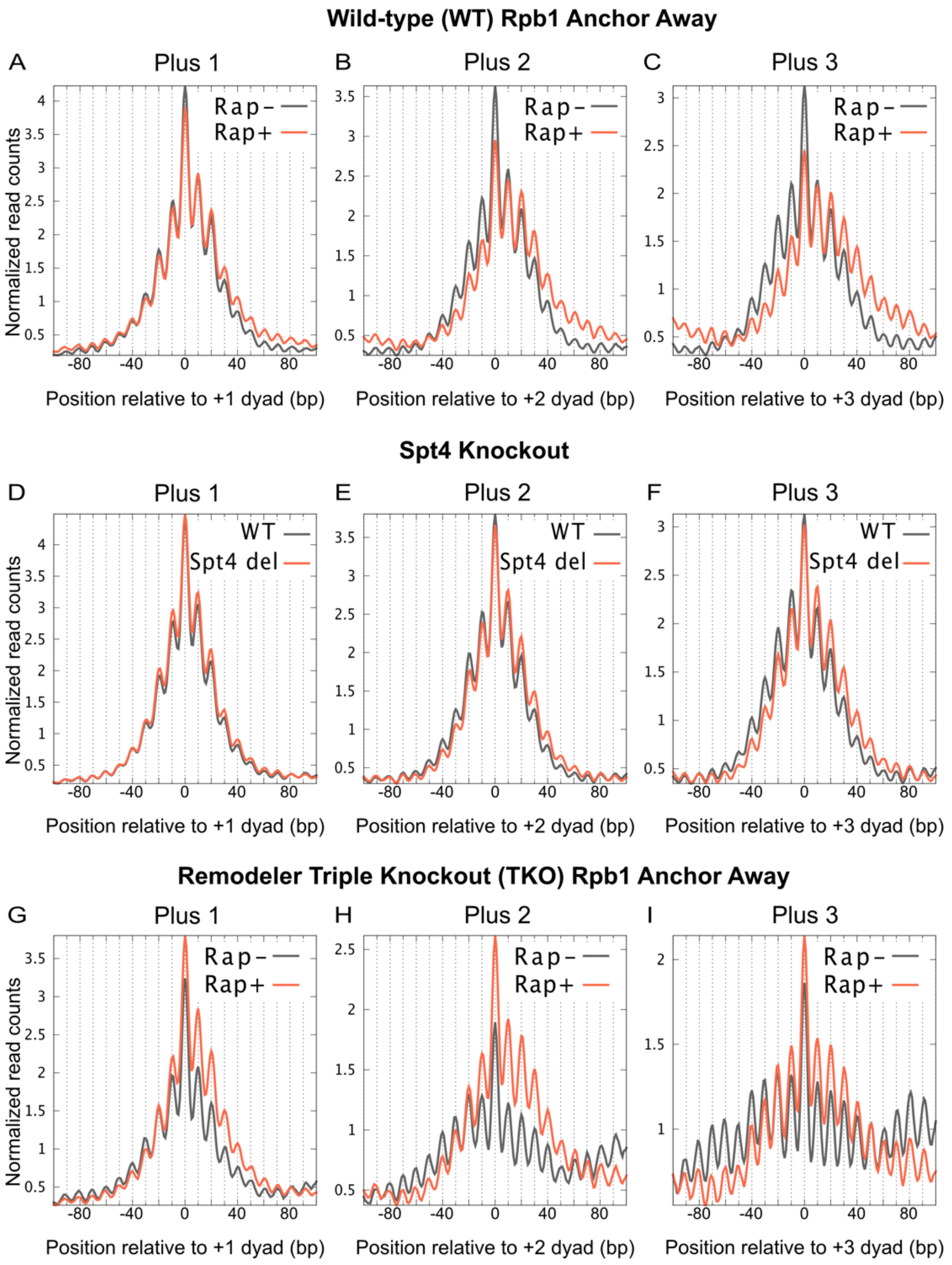


**Figure S7. Nucleosome shifts due to perturbations in transcription are also quantized.** High-resolution plot of MNase-seq data at TSS+1, TSS+2, and TSS+3 nucleosomes for the indicated strains and perturbations.

**Figure S8**

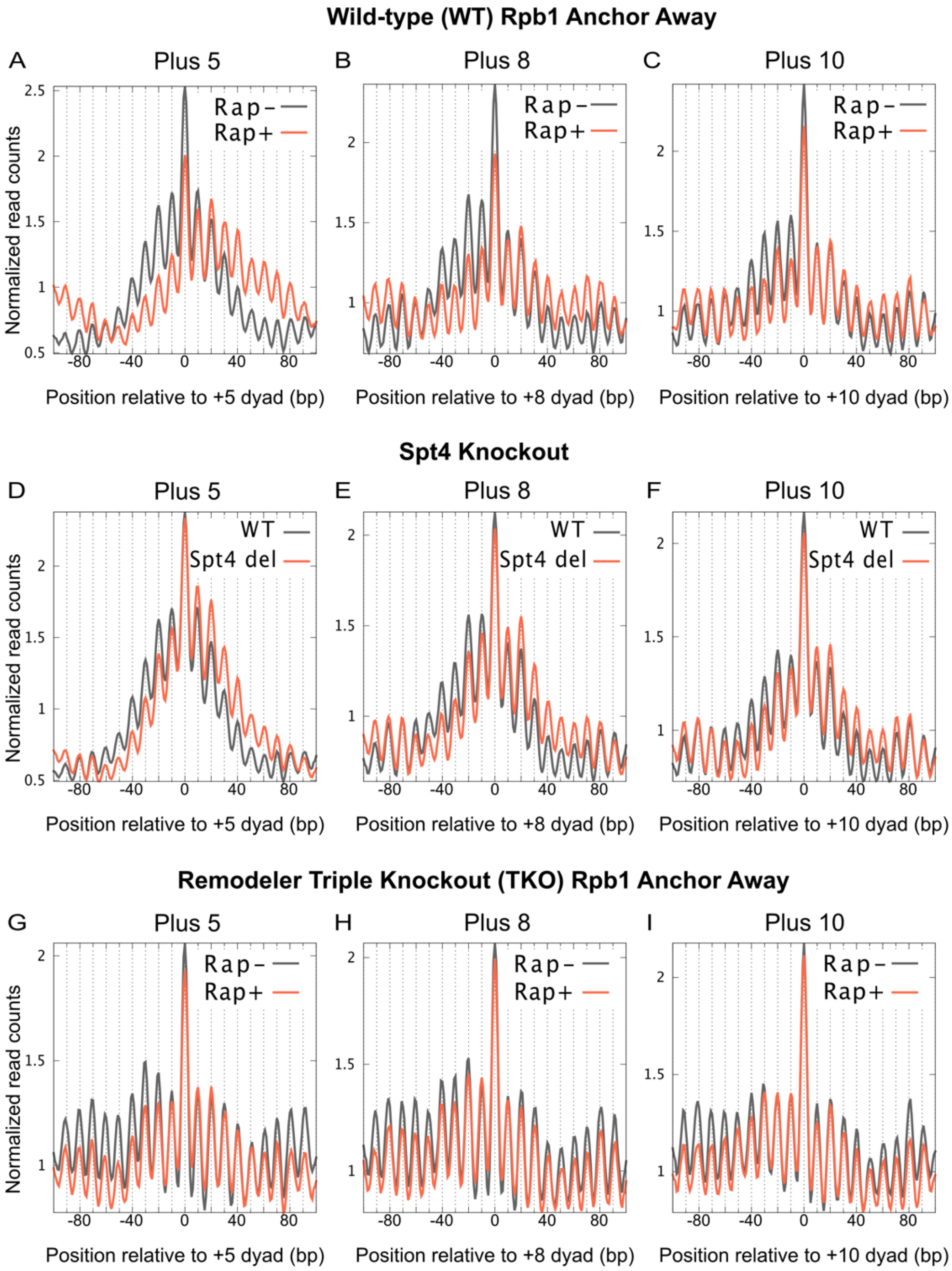


**Figure S8. Nucleosome shifts due to perturbations in transcription are also quantized.** High-resolution plot of MNase-seq data at TSS+5, TSS+8, and TSS+10 nucleosomes for the indicated strains and perturbations.
